# Interdependent mode of RNA recognition by the interacting KH-domain proteins KhpA and KhpB

**DOI:** 10.64898/2026.10.12.757534

**Authors:** Manasa Narayan, Sahil Sharma, Thorsten Bischler, Nay Won Lett, Tom Gräfenhan, Sander Granneman, Katherine E. Berry, Mona Alzheimer, Cynthia M. Sharma

## Abstract

RNA-binding proteins (RBPs) are central regulators of gene expression. In bacteria, two KH domain-containing proteins, KhpA and KhpB, are increasingly recognized as globally-acting RBPs, including in pathogens that lack well-studied RBPs, such as Hfq and ProQ. While KhpA and KhpB interact with various RNAs in diverse bacteria, their crosstalk and molecular basis of RNA recognition remain poorly understood. Here, we explored the mutual dependence between KhpA and KhpB in the food-borne pathogen *Campylobacter jejuni*. We demonstrate KhpA and KhpB interact *in vivo*, and their individual global RNA recognition is largely dependent on the presence of the partner protein. Using RNA co-immunoprecipitation, we reveal that KhpA requires a conserved KH-domain motif for RNA binding, whereas KhpB relies on an atypical R3H domain. Moreover, mutating the RNA-binding domain of one protein compromises the RNA interactome of the other. Overall, we demonstrate an interdependent RNA-binding mode of KhpA and KhpB and anticipate that such KH-proteins have co-evolved to elicit joint RNA regulation in bacteria.

## Introduction

In all kingdoms of life, as well as in viruses, RNA-binding proteins (RBPs) are central players in RNA metabolism and gene regulation, controlling RNA stability, translation, and localization. RBPs typically recognize their RNA ligands through distinct RNA-binding domains (RBDs), including widespread structural motifs, such as the K-homology (KH) domain, RNA recognition motif (RRM), and the S1 domain (1). Much of our current understanding of RBDs originates from studies in eukaryotes, where multiple RBDs within a single protein or across several proteins often cooperate to enhance RNA binding (2–4). In contrast, little is known about such crosstalks in bacterial RBPs, which are generally more streamlined, containing fewer RBDs within a protein.

In bacteria, RBPs are involved in shaping microbial fitness and virulence-related processes at transcriptional, post-transcriptional, and translational levels (5–8). CsrA, Hfq, and ProQ are the best-characterized globally-acting RBPs, which mediate post-transcriptional control in diverse bacteria. However, many bacteria lack homologs of these proteins, raising the need to understand how RNA-mediated regulation operates across diverse bacterial lineages. A growing body of recent research has highlighted a pair of KH-domain containing proteins, KhpA and KhpB, as candidate RBPs in diverse species (9–13). While KhpA and KhpB are broadly conserved among bacterial species and are emergingly recognized for their potential global RNA-binding roles (14), their functions and mechanisms are still unclear. Previous work in *Streptococcus pneumoniae* suggested that KhpA and KhpB interact with each other. Moreover, despite being encoded in distant genomic loci, *khpA* and *khpB* often co-occur (14) and have been implicated in various cellular processes, including cell elongation, toxin production, and ethanolamine utilization in various bacteria (9, 10, 12, 15, 16). Both *khpA* and *khpB* exhibit conserved gene synteny within housekeeping gene operons alongside genes involved in tRNA maturation, ribosome assembly, and protein insertion into membranes, indicating some conserved cellular function (14).

The global RNA-binding capacities of KhpA and/or KhpB were mainly uncovered through RNA co-immunoprecipitation experiments, which suggested potential associations with diverse transcripts, including mRNAs and small RNAs (sRNAs) (9–12). In species such as *S. pneumoniae* and *F. nucleatum,* where both KH-proteins have been studied, they seem to share overlapping RNA targets (9, 12). The genome of the Gram-negative Epsilonproteobacterium, and currently most-common cause of bacterial food-poisoning *Campylobacter jejuni* (17, 18), lacks homologs of Hfq and ProQ but encodes homologs of KhpB (*Cj0957c*) and KhpA (*Cj0711*) (19). In our lab, we recently also captured KhpB in a global screen for RBPs in *C. jejuni* and observed a similar overlap in the RNA targetome of KhpA and KhpB in *C. jejuni* in RIP-seq experiments (Sharma Lab, unpublished). The observed pattern of overlapping targets of KhpA/B across species suggests an interdependent RNA-binding model between KhpA and KhpB, reminiscent of KH domain proteins in eukaryotes (20). Nonetheless, the extent to which these proteins functionally depend on each other, especially in terms of RNA interactions, as well as their molecular determinants of RNA interaction, remain unknown.

While both proteins feature a single KH domain, KhpB harbors another RBD, an R3H domain, at the C-terminus, and an N-terminal Jag-N domain of unknown function ((14) and Fig. 1A). KH domains are highly conserved RNA-binding motifs characterized by a canonical GXXG hydrophobic groove that mediates nucleic acid interactions (20–22), whereas R3H domains likely recognize single-stranded nucleic acids via an RXXXH motif (23). While the overall domain architecture of KhpA and KhpB is conserved across bacterial homologs, the proteins show considerable amino acid sequence diversity (14). The GXXG and RXXXH RNA-binding motifs are generally preserved among other studied KhpA/B homologs. In *C. jejuni*, KhpA also has an intact GXXG motif, whereas KhpB has a mutated GXXG motif (GKEA) and its R3H domain lacks the RXXXH motif (Fig. 1A; Fig. S1). Which domains mediate RNA binding in each protein is still largely unknown. Moreover, as no *in-vivo* data currently define the molecular basis of target RNA recognition by KhpA and KhpB, the functional implications of the different RBDs for target binding remain largely unknown.

**Fig. 1:**
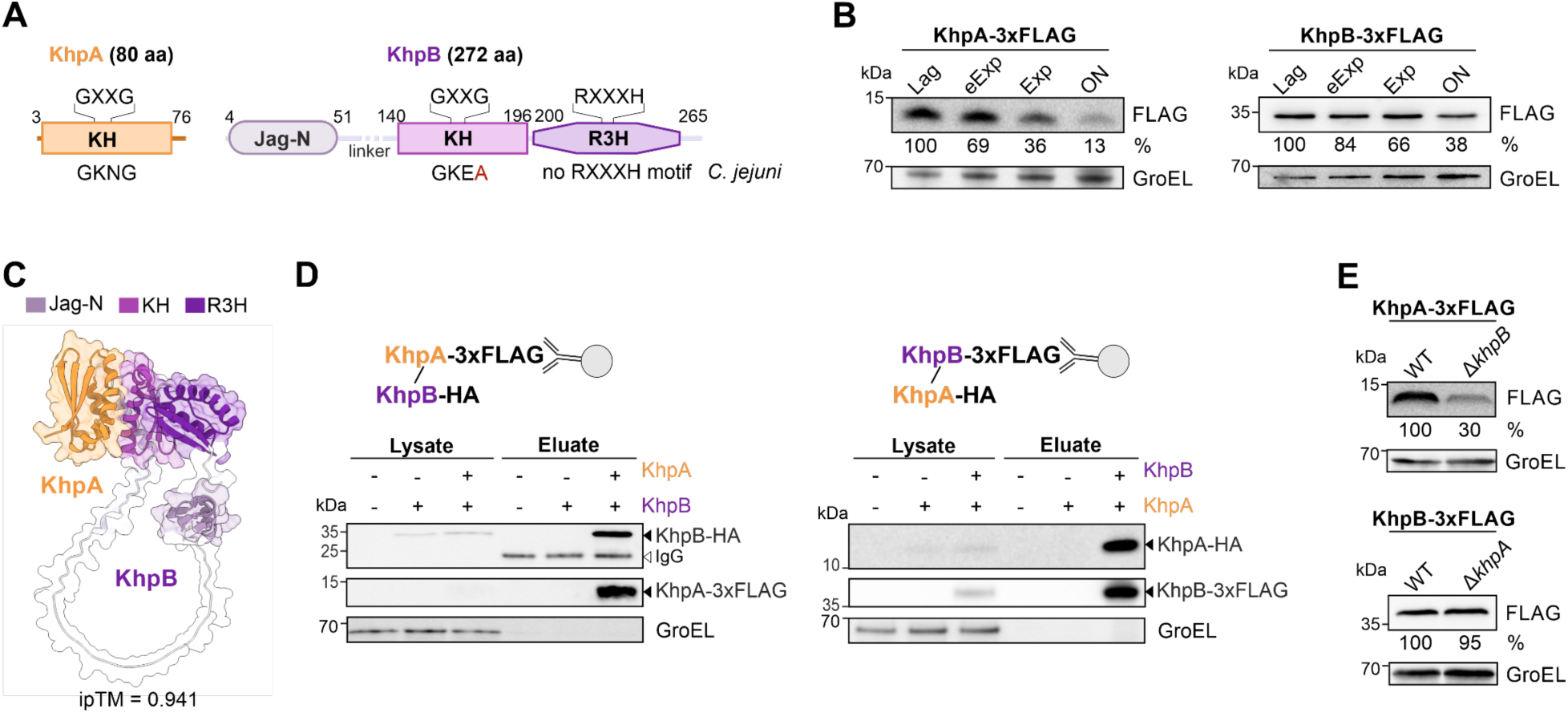
Domain architecture, expression, and interaction analyses of *C. jejuni* KhpA and KhpB. **(A)** Schematic of the domain architecture and conserved residues of KhpA and KhpB in *C. jejuni.* Red: degenerate amino acid in the characteristic GXXG motif. **(B)** Western blot analysis of 3xFLAG-tagged KhpA and KhpB in *C. jejuni* detected with an anti-FLAG antibody during growth. GroEL served as a loading control. Bands were quantified using ImageJ (61), normalized to GroEL, and percentages relative to the lag phase are given. eExp: early exponential. ON: overnight. Blots are representative of at least three biological replicates. **(C)** Predicted heterodimer structure of *C. jejuni* KhpA and KhpB using AlphaFold2-multimer (24). Domain colours correspond to those in (A). Interface predicted template modelling (ipTM), denoting the model confidence, is shown below. **(D)** Reciprocal co-immunoprecipitation of KhpA-/KhpB-3xFLAG from *C. jejuni* lysates. 3xFLAG fusions were immunoprecipitated and detected on western blots with an anti-FLAG antibody. Co-purified KhpB-/KhpA-HA were detected using an anti-HA antibody. GroEL served as a loading control. -/- denotes the untagged WT control. + indicates the presence of the epitope-tagged gene at the native locus. Representative blots of at least two biological replicates are shown. **(E)** Western blot of KhpA-3xFLAG and KhpB-3xFLAG expression in *C. jejuni* WT and the deletion mutant of *khpB* and *khpA*, respectively. FLAG-tagged proteins were detected with an anti-FLAG antibody, and GroEL served as a loading control. Bands were quantified as in (A). Blots are indicative of at least three independent replicates.

In this study, we dissect the *in-vivo* molecular determinants of KhpA and KhpB for RNA-binding and demonstrate their mutual functional dependence in *C. jejuni*. We show that KhpA and KhpB interact with each other, and the *in-vivo* RNA interactome of each protein depends on the presence of the interacting partner. Mutagenesis analysis combined with global RIP-seq analyses (RNA co-immunoprecipitation combined with sequencing) revealed that the conserved KH domain of KhpA is essential for nearly all of its RNA interactions. In contrast, for KhpB, it is the R3H domain, despite lacking the canonical RXXXH motif, that plays the key role in RNA binding, rather than the KH domain or the Jag-N domain. Moreover, we demonstrate that KhpA and KhpB function through their mutually dependent RNA-binding properties, and propose that these KH domain proteins may have co-evolved to fulfill their joint role in bacterial RNA regulation.

## Results

### KhpA and KhpB interact in *C. jejuni*, with KhpB impacting KhpA protein levels

To explore functional connections between KhpA and KhpB, we first examined their expression during different growth phases in *C. jejuni*. Both KhpA and KhpB are expressed throughout the growth of *C. jejuni*, though levels of both proteins decrease upon entry into the stationary phase (Fig. 1B). Using AlphaFold2-multimer structural predictions (24), we next examined a potential interaction of the two proteins and detected a high-confidence interaction between the KH domains of *C. jejuni* KhpA and KhpB (Fig. 1C). To further experimentally validate the KhpA/B interaction *in vivo*, we first tested their interaction in a bacterial two-hybrid (B2H) system (Fig. S2). This revealed a robust interaction between C. jejuni KhpA and KhpB. This observation is consistent with a previous yeast two-hybrid screen that supported an interaction between the two *C. jejuni* proteins (25) as well as studies supporting interaction of *S. pneumoniae* KhpA and KhpB (9, 16). Next, to confirm interaction in the native host, we generated a *C. jejuni* strain expressing KhpA-3xFLAG and KhpB-HA from their endogenous loci and performed co-immunoprecipitation (co-IP) using an anti-FLAG antibody (Fig. 1D). Western blot analysis confirmed the successful pull-down of KhpA-3xFLAG and co-purification of KhpB-HA, supporting an *in-vivo* interaction between the two proteins in *C. jejuni*. Reciprocally, KhpA-HA could be successfully co-purified with KhpB-3xFLAG. After demonstrating that the two proteins interact *in vivo*, we next investigated whether the expression of KhpA and KhpB depends on the presence of their respective interaction partner. While KhpB expression remained unchanged in the absence of *khpA*, KhpA protein levels decreased nearly threefold upon deletion of *khpB* in *C. jejuni* (Fig. 1E). Overall, the expression and co-IP analyses show that KhpB interacts with KhpA and impacts its steady-state protein levels.

### The RNA targetome of KhpA and KhpB depends on the presence of the interacting partner

Since KhpA and KhpB interact at the protein level, we reasoned that their RNA-binding capacity and resulting RNA targetomes may depend on the presence of the respective interacting partner RBP. To explore this, we performed RIP-seq using C-terminally FLAG-tagged KhpA and KhpB in the wild-type (WT) and respective Δ*khpB* / Δ*khpA* partner protein deletion background (Fig. 2A). Untagged WT and deletion strains served as respective background controls. Despite reduced KhpA levels in Δ*khpB*, we successfully recovered both proteins in co-IP fractions of WT and Δ partner background strains (Fig. S3). Principal component analysis (PCA) of the RIP-seq libraries generated from co-purified RNAs revealed that the KhpA-3xFLAG and KhpB-3xFLAG samples in the WT background clustered separately from one another and from the untagged WT control, indicating distinct RNA-binding profiles for both proteins (Fig. S4A). In contrast, in the absence of their respective partner (Δ*khpB* for KhpA and Δ*khpA* for KhpB), KhpA- and KhpB-bound RNA profiles clustered closer to the untagged libraries, indicating loss of target enrichment. Consistent with this, RIP-seq of KhpA-3xFLAG and KhpB-3xFLAG in the WT background showed strong enrichments of CDSs and 5’UTRs (Fig. 2B; Table S1). This enrichment was clearly lost in the Δ partner backgrounds, reinforcing that KhpA and KhpB require each other for target enrichment.

**Fig. 2:**
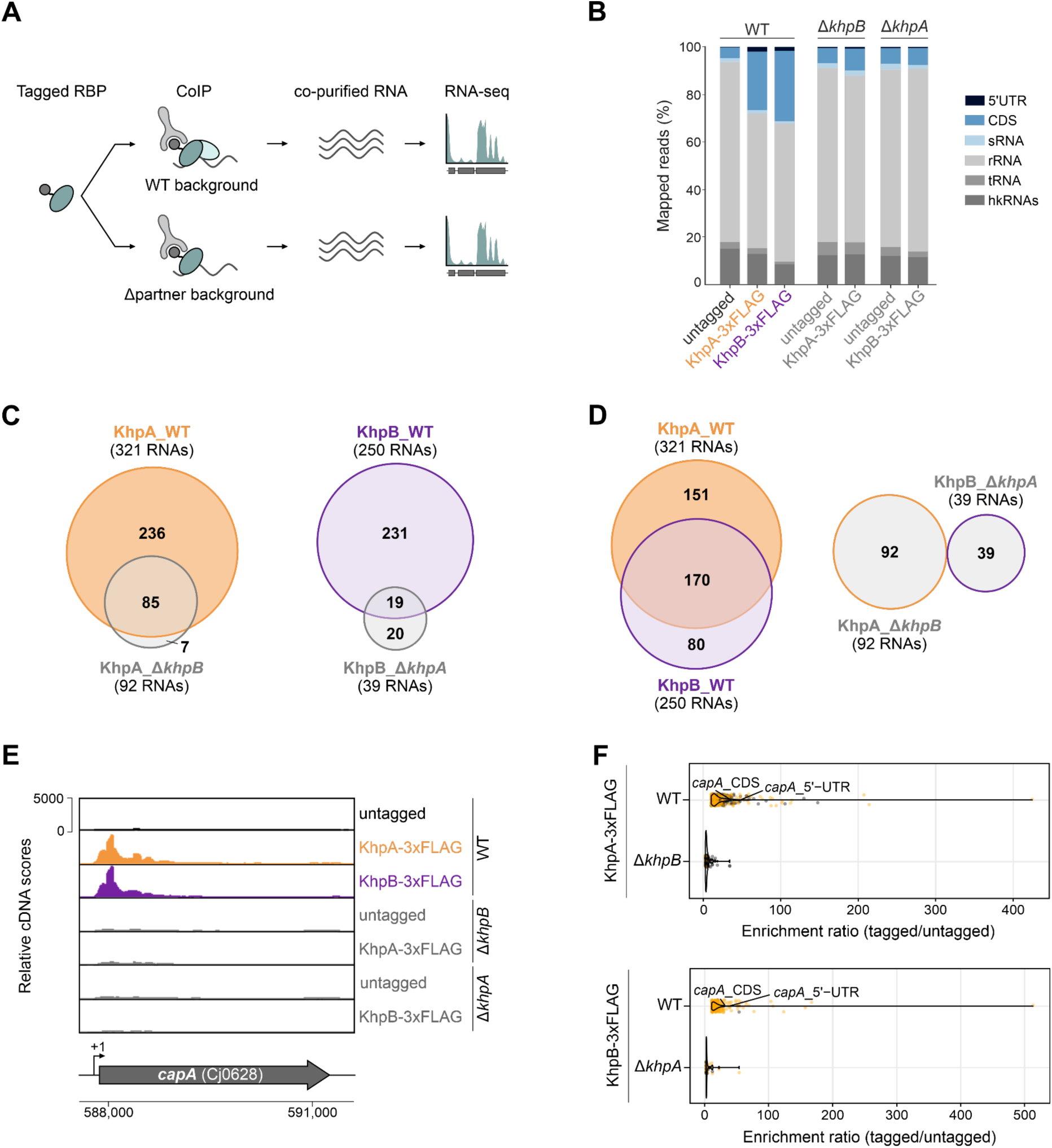
RIP-seq analyses of *C. jejuni* KhpA and KhpB in the absence of the interacting partner protein. **(A)** Workflow depicting the co-immunoprecipitation (co-IP) and sequencing of RNAs bound by FLAG-tagged RBP in the presence (WT background) or absence (Δ partner background) of the protein interaction partner. **(B)** Bar graphs showing the mapped read distribution percentage of the indicated RNA classes in the untagged, KhpA-3xFLAG, and KhpB-3xFLAG co-IP libraries in the WT or deletion background strains of the partner. **(C)** (Left) Overlap between the RNA targetome of KhpA-3xFLAG in WT (orange) versus Δ*khpB* (grey). (Right) Overlap between the RNA targetome of KhpB-3xFLAG in WT (purple) versus Δ*khpA* (grey). Numbers indicate RNAs with a log_2_ fold change ≥ 1, *p*-adj ≤ 0.05, and an average CPM (counts per million) ≥10 in 3xFLAG libraries. **(D)** Comparison of the RNA targetomes of KhpA-3xFLAG and KhpB-3xFLAG pull-downs in WT (left) and Δ partner (right) backgrounds. Criteria for the numbers indicated are the same as in (C). **(E)** Genome browser view depicting cDNA read coverage of co-IP libraries mapped to the *Campylobacter* adhesion protein (*capA*) locus in the WT (upper) and deletion background of the partner (lower). One representative biological replicate of untagged controls, KhpA-3xFLAG, and KhpB-3xFLAG in WT (black, orange, purple, respectively) and Δ partner (gray) backgrounds is shown. The transcriptional start site (TSS) is denoted as a black arrow (58). **(F)** (Top) Violin plot depicting the RIP-seq gene enrichment ratio of KhpA-3xFLAG in WT versus Δ*khpB.* (Bottom) Violin plot depicting the RIP-seq gene enrichment ratio of KhpB-3xFLAG in WT versus Δ*khpA*. Unique and commonly enriched RNAs between two backgrounds are represented by yellow and gray points, respectively. The enrichment ratio for each significantly enriched gene is calculated by dividing the average CPM of the tagged library by that of the untagged library.

DESeq2 analysis of significantly enriched transcripts in the WT background revealed 321 and 250 transcripts co-purified with KhpA and KhpB, respectively (log_2_ fold-change ≥ 1, *p*-adj ≤ 0.05; average counts per million ≥ 10 in tagged libraries). In contrast, deletion of the partner KH-protein resulted in a drastic reduction of co-purified RNAs by ∼71% and ∼84% for KhpA and KhpB (92 for KhpA, 39 for KhpB), respectively (Fig. 2C; Table S2). Additionally, both KhpA and KhpB enriched 7 and 20 unique targets in the absence of the partner protein, suggesting possible partner-dependent specificity of targets (Fig. 2C and Fig. S4B). While KhpA and KhpB share a largely overlapping RNA targetome in the WT background, this overlap is lost in the absence of the partner protein (Fig. 2D). For example, among the top significantly enriched KhpA/B targets in the WT background is the mRNA encoding *Campylobacter* adhesion protein (CapA) (Fig. S4C). While *capA* is enriched in the RIP-seq of both KhpA/B, no enrichment is visible in the Δ partner backgrounds (Fig. 2E). Several other commonly enriched targets were also de-enriched in Δ partner backgrounds (Fig. S4C). Moreover, several KhpA-specific targets (e.g., *ksgA, flhB*, *pyrF*) were among the top targets for KhpA in the Δ*khpB* background (Fig. S4C). In the Δ*khpA* background, several unique targets that were not enriched in the KhpA/B WT background emerged as top targets for KhpB (Fig. S4D). Among such transcripts still bound in the partner deletion mutant, the overall enrichment ratio was strongly reduced for both KhpA and KhpB in the absence of the partner (Fig. 2F), indicating that the partner protein also influences target RNA enrichment. Together, our findings demonstrate that the presence of both KH-proteins is crucial for mediating the majority of their RNA interactions in *C. jejuni*.

### The KH domain of KhpA and the R3H domain of KhpB are the RNA-binding determinants for most bound RNAs

To investigate how the different domains and potential RNA binding motifs of KhpA/B contribute to their RNA-binding capacities, we applied the pyRBDome pipeline (26) to predict potential RNA-binding amino acid residues in *C. jejuni* KhpA and KhpB. For comparison, we also included the KhpA/B homologs from *S. pneumoniae*, *C. difficile,* and the related gastric pathogen *Helicobacter pylori*. Briefly, the pyRBDome pipeline submits Alphafold models to five independent protein-RNA (and protein-ligand) binding site prediction algorithms that use sequence, structure or both to identify putative RNA-binding residues. An XGBoost ensemble model then uses the outputs of the individual predictors to make a final prediction. This revealed high RNA-binding probabilities for many amino acids in all KhpA homologs analysed, including *C. jejuni* KhpA, with their conserved GXXG motif being the most likely RNA-binding site (Fig. S5-S6). KhpB homologs showed variable RNA-binding scores, with *C. difficile* KhpB ranking the highest and *C. jejuni* KhpB the lowest (Fig. S7-S8). The conserved GXXG motif was one of the likely RNA-binding regions in all KhpB homologs, except for *C. jejuni* KhpB, which showed the R3H domain as the most probable RNA-binding interface.

To validate these predictions for *C. jejuni* KhpA/B *in vivo*, we complemented the respective deletion backgrounds with KhpA/B variants that carry point mutations or domain truncations at the native locus (Fig. 3A). The characteristic GXXG motif of the KH domain can be disrupted by replacing residues to modulate the net charge on the nucleic acid-binding surface (27). Accordingly, we mutated the KhpA KH motif from GKNG to GDDG (KhpA_KH^GDDG^) and that of KhpB from GKEA to GDDA (KhpB_KH^GDDA^). In addition, we generated truncated mutants of KhpB by individually deleting the Jag-N domain (KhpB_ΔJag) and the non-canonical R3H domain (KhpB_ΔR3H). All constructs were also 3xFLAG-tagged to facilitate protein detection, and all mutants expressing these constructs showed similar growth to WT under the examined conditions (Fig. S9). While protein levels of KhpB mutant variants were comparable to WT, KhpA_KH^GDDG^ showed reduced expression (Fig. 3B).

**Fig. 3:**
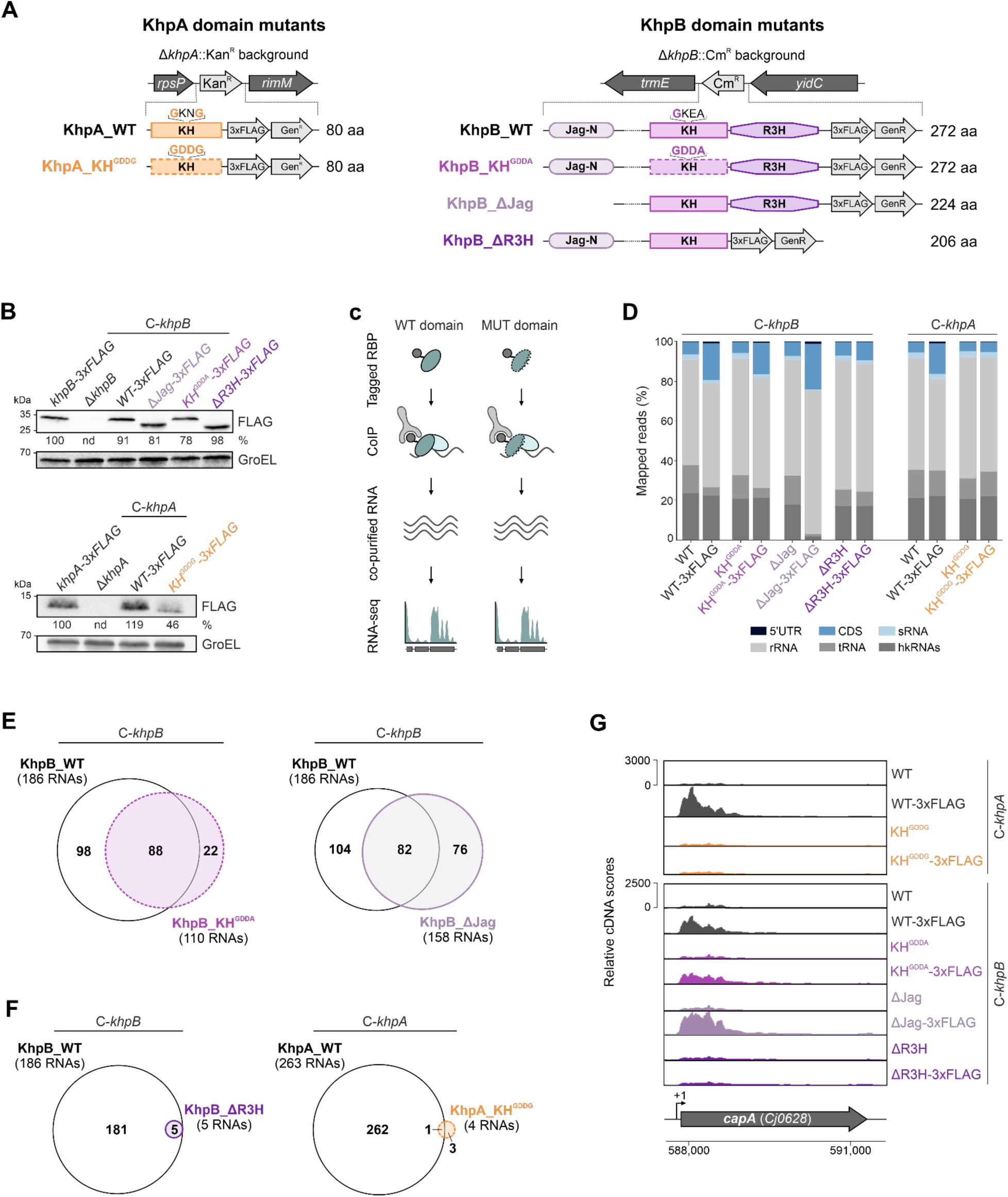
RIP-seq analyses of domain and point mutants of KhpA and KhpB in *C. jejuni*. **(A)** (Left) Schematic representation of the complementation of WT and GXXG mutant variants of the KH-domain of KhpA at the native *khpA* locus of Δ*khpA.* (Right) Schematic representation of the complementation of WT and different domain mutant variants of KhpB at the native *khpB* locus of Δ*khpB.* All complemented variants include a C-terminal 3xFLAG-tag and a non-polar gentamicin resistance marker. **(B)** Western blot analysis of FLAG-tagged WT and domain mutant variants from (A), detected using an anti-FLAG antibody. Bands were quantified using ImageJ, normalized to GroEL, and percentages relative to KhpA-3xFLAG in *C. jejuni* WT (upper) and KhpB-3xFLAG (lower) in *C. jejuni* WT are given. Blots are representative of at least two biological replicates. **(C)** RIP-seq workflow of WT and domain mutant variants of KhpA-3xFLAG and KhpB-3xFLAG from (A). **(D)** Bar graphs showing the mapped read distribution percentage of the indicated RNA classes in the WT and domain mutant variants of KhpA-3xFLAG and KhpB-3xFLAG co-IP libraries. Each FLAG-tagged strain is compared to its respective untagged strain. **(E)** Overlap between the RNA targetome of WT and domain mutant variants of KhpB-3xFLAG that retain RNA-binding ability. (Left) Venn diagram comparing functional (WT) KhpB-3xFLAG (purple) versus Jag-N truncated KhpB-3xFLAG (gray). (Right) Venn diagram comparing functional (WT) KhpB-3xFLAG (purple) versus GXXG mutated KhpB-3xFLAG (pink). Numbers indicate RNAs with log_2_ fold-change ≥1, *p*-adj≤0.05, and average CPM ≥10 in tagged libraries. **(F)** Overlap between the RNA targetome of WT and RNA-binding mutant variants of KhpA-3xFLAG and KhpB-3xFLAG. (Left) Venn diagram comparing functional (WT) KhpB-3xFLAG (purple) versus R3H truncated KhpB-3xFLAG (maroon). (Right) Venn diagram comparing functional (WT) KhpA-3xFLAG (orange) versus GXXG mutated KhpA-3xFLAG (red). The criteria for numbers indicated are the same as in (E). **(G)** cDNA read coverage for the functional (WT) and domain mutant variants of KhpA-3xFLAG and KhpB-3xFLAG co-IP libraries mapped to the *Campylobacter* adhesion protein (*capA*). Each FLAG-tagged strain is compared to its respective untagged strain. TSS is denoted as a black arrow.

To determine the impact of the domain mutations on global RNA-binding, we performed RIP-seq of FLAG-tagged WT and KhpA/B domain mutant variants (Fig. 3C). Untagged WT or mutant variants served as controls (Fig. S10A), and western blot analysis of co-IP fractions showed specific recovery of WT and domain mutants of KhpA/B (Fig. S10B-D). PCA plots of the resulting RIP-seq libraries of copurified transcripts showed a clear separation of KhpA_WT-3xFLAG, KhpB_WT-3xFLAG, KhpB_ΔJag-3xFLAG, and KhpB_KH^GDDA^-3xFLAG from their respective untagged libraries, highlighting distinct RNA profiles of these mutants (Fig. S11A). In contrast, KhpA_KH^GDDG^-3xFLAG and KhpB_ΔR3H-3xFLAG co-clustered with their untagged matching controls, strongly indicating a loss of target enrichment specific to these mutants. Supporting this, RIP-seq reads showed a clear enrichment of CDSs in KhpB_WT-3xFLAG, KhpB_KH^GDDA^-3xFLAG, and KhpB_ΔJag-3xFLAG as well as in KhpA_WT-3xFLAG libraries as compared to respective untagged controls (Fig. 3D; Table S1). This enrichment was strikingly lost in KhpA_KH^GDDG^-3xFLAG and KhpB_ΔR3H-3xFLAG libraries (Fig. 3D), suggesting a crucial role for the canonical KH domain of KhpA and the degenerate R3H domain of KhpB in RNA binding.

Next, we looked at the overall number of enriched RNAs for each of the complemented mutant protein variants above. We identified 263 and 186 significantly enriched RNAs for KhpA_WT and KhpB_WT, respectively, indicating that complementation of the deletion mutants with WT versions of KhpA/B (cKhpA-WT and cKhpB-WT) restores global protein-RNA interactions similar to those observed during RIP-seq with tagged KhpA/B in WT (Fig. 3E and Fig. S11B; Table S3). As expected, KhpB_ΔJag and KhpB_KH^GDDA^ also retained RNA-binding capacities, enriching 158 and 110 transcripts, respectively, both of which largely overlapped with those co-purified with KhpB_WT (Fig. 3E). This indicates that the diverged GKEA motif of KhpB is not strictly required for its direct RNA interaction. Although KhpB_ΔJag-3xFLAG libraries had lower total read counts and reduced tRNA and housekeeping RNA reads, the enrichment of target CDSs remained prominent (Fig. 3D). Of note, KhpB_ΔJag also enriched around 45 unique non-KhpA/B targets (Fig. S11B), suggesting that the Jag-N domain offers additional specificity. Strikingly, KhpA_KH^GDDG^ and KhpB_ΔR3H enriched only four and five transcripts, which partially or fully overlapped with KhpA_WT and KhpB_WT, respectively (Fig. 3F; Fig. S11B). This demonstrates that the KH domain of KhpA and the R3H domain of KhpB are indispensable for RNA interaction. Moreover, top targets of KhpA, including KhpA-specific targets, were not enriched in KhpA_KH^GDDG^, and those of KhpB were not significantly enriched in KhpB_ΔR3H (Fig. S11C). Reads mapping to *capA* mRNA also showed enrichment in all WT and domain mutant variants of KhpA and KhpB, whereas this enrichment was lost in KhpA_KH^GDDG^ and KhpB_ΔR3H (Fig. 3G). Overall, these results demonstrate that, while the KhpB KH motif and the Jag-N domain are not vital for its RNA binding, the conserved GXXG motif of KhpA and the R3H domain without the RXXXH motif of KhpB are the essential RNA-binding domains of the two KH-proteins in *C. jejuni*.

### KhpA and KhpB interact in an RNA-independent manner

So far, it is unclear whether KhpA and KhpB require RNA-binding to interact with each other or whether they can form a complex independent of RNA. To explore this, we next investigated whether the RNA-binding deficient mutants could still bind their partner KH-protein or if their loss of RNA binding results in disrupted partner interaction. Alphafold2-multimer structural predictions suggested the KH domain and IDR (intrinsically disordered region) of KhpB may be contact points with KhpA (Fig. 1C). Supporting this, Alphafold2-multimer predicted a high-confidence interaction between all tested domain mutants and functional partner protein (Fig. S12). To experimentally validate the interaction *in vivo*, we performed co-IP experiments using anti-FLAG antibodies on strains expressing KhpA-3xFLAG, along with KhpB-HA, both WT and mutant versions, from their native loci. Western blot analysis confirmed that all KhpB domain and point mutants tested retained their ability to interact with KhpA (Fig. 4A). Similarly, an anti-FLAG co-IP in a strain expressing KhpA_KH^GDDG^-3xFLAG together with KhpB-HA confirmed that RNA-binding-deficient KhpA still interacts with wild-type KhpB (Fig. 4B). Since KhpA protein levels decreased in the absence of KhpB (Fig. 1E), we tested whether RNA-binding-deficient KhpB or other domain mutants affected KhpA expression levels. None of the domain mutants altered KhpA protein levels compared to WT, suggesting that the interaction between KhpA and the tested domain mutants of KhpB maintains KhpA steady-state levels (Fig. S13).

**Fig. 4:**
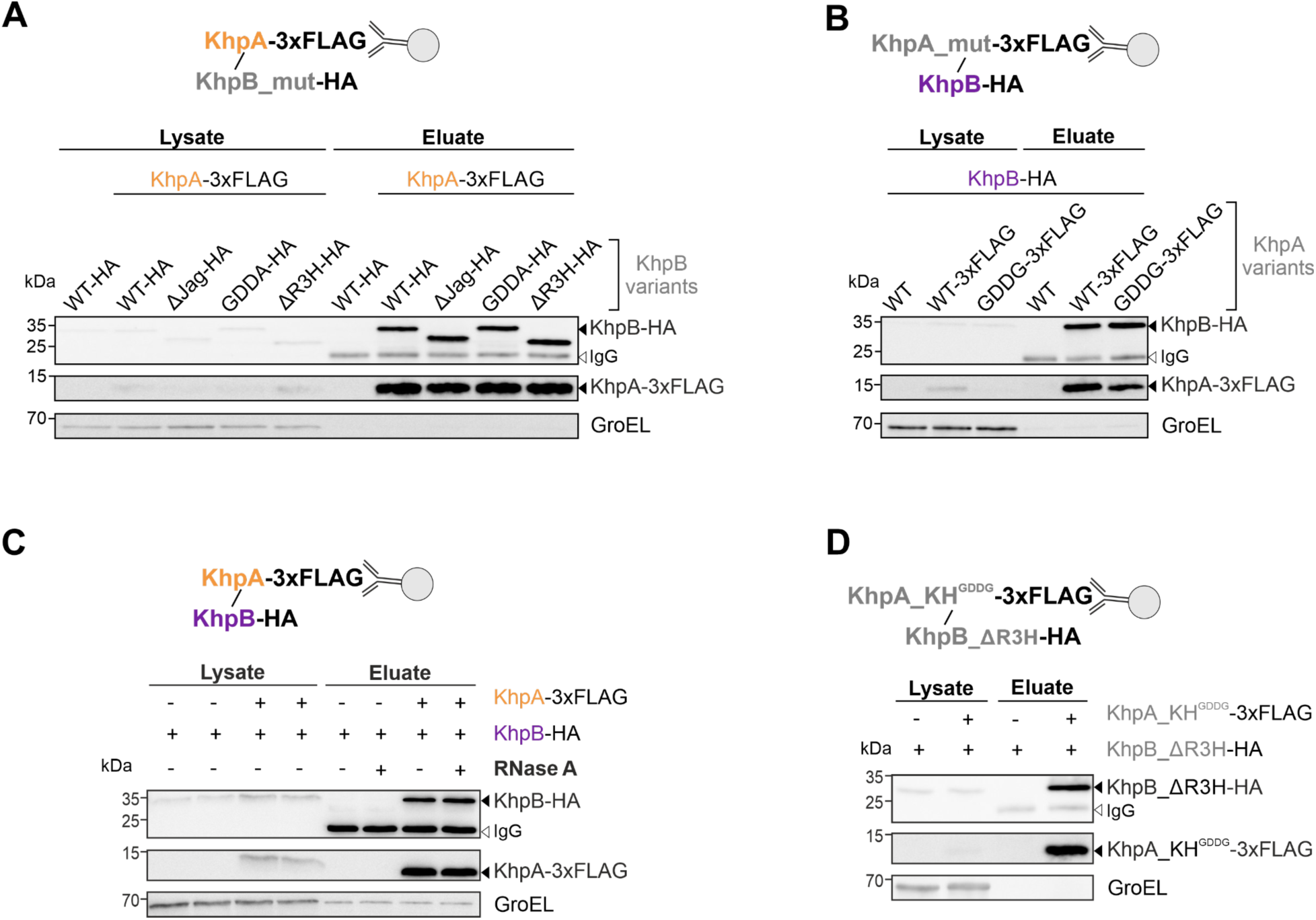
Protein interaction analyses of *C. jejuni* KhpA and KhpB domain and point mutants. **(A)** Co-immunoprecipitation (co-IP) of KhpA-3xFLAG, combined with western blot analysis. **(B)** Co-IP of WT and domain mutant variants of KhpA-3xFLAG, combined with western blot analysis. **(C)** Co-IP of KhpA-3xFLAG after with (+) or without (-) RNase A treatment, combined with western blot analysis. **(D)** Co-IP of GDDG-mutated KhpA-3xFLAG, combined with western blot analysis. **(A-D)** 3xFLAG fusions were immunoprecipitated and detected on western blots with an anti-FLAG antibody. Co-purified KhpB-HA variants were detected using an anti-HA antibody. GroEL served as a loading control. + indicates the presence of the epitope-tagged gene at the native locus. Representative blots of at least two biological replicates are shown.

Next, we further assessed whether the RNA interaction of both KhpA and KhpB is necessary to mediate their protein-protein interaction. Therefore, we treated exponential-phase lysates from the KhpA-3xFLAG and KhpB-HA dual-tagged strain with RNase A, followed by co-IP and western blot analysis. Pull-down of KhpA-3xFLAG post RNase A treatment co-purified KhpB-HA, indicating that their interaction is not sensitive to RNase treatment (Fig. 4C). To further dissect the role of RNA in this interaction, we performed co-IP on a strain expressing RBD mutants of both proteins, KhpB_ΔR3H-HA and KhpA_KH^GDDG^-3xFLAG (Fig. 4D). Both RBD mutants retained their interaction, suggesting that neither the KhpB R3H-domain nor the KhpA KH-motif is involved in their direct interaction, nor that their binding to RNA is required for KhpA-KhpB complex formation. Together, our data strongly suggest that direct interaction between the two KH domain proteins does not involve RNA.

### RNA-binding-deficient Khp partner protein compromises the RNA targetome of the KhpA/B heterocomplex

Our RIP-seq experiments demonstrated that the absence of one partner KH protein drastically reduces the RNA-binding activity of the other (Fig. 2). Therefore, we hypothesized that the presence of an RNA-binding-deficient partner could also influence the RNA-binding ability of the functional counterpart. To address this, we performed RIP-seq of a WT protein in the presence of an RNA-binding-deficient partner protein: KhpA-3xFLAG in *khpB*_WT and *khpB*_ΔR3H backgrounds, and KhpB-3xFLAG in *khpA*_WT and *khpA_*KH^GDDG^ backgrounds were subjected to co-IP (Fig. 5A, Fig. S14A). Respective untagged background controls were included, and successful pull-down of FLAG-tagged proteins was confirmed on western blots (Fig. S14B). PCA of RIP-seq libraries in the presence of WT partner showed distinct clustering of KhpA-3xFLAG and KhpB-3xFLAG from untagged controls, indicating specific RNA enrichments as expected (Fig. S15A). This separation was almost lost in the RBD mutant partner background (PC1, 44% variance), but partial retention on PC2 (11% variance) suggests residual RNA binding. In support of this, while the KhpA and KhpB pull-downs in the WT partner background showed the expected enrichment of 5’ UTRs and CDSs, this enrichment was notably reduced in the RBD mutant background (Fig. 5B; Table S1), highlighting the impact of the partner RBD mutation on the RNA targetome of the functional other.

**Fig. 5:**
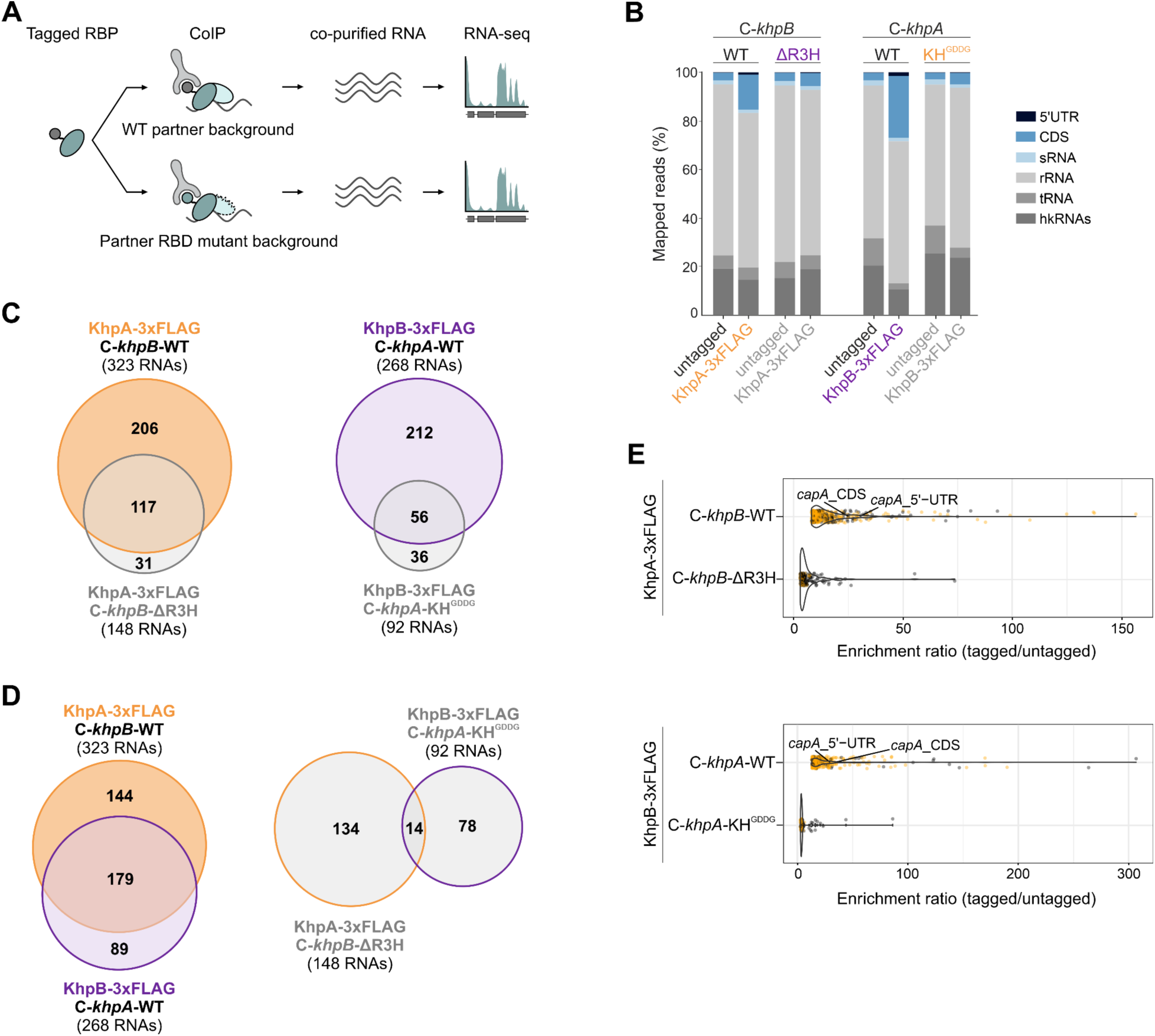
RIP-seq analyses of *C. jejuni* KhpA and KhpB in the presence of RNA-binding-deficient partner protein. **(A)** Workflow describing the co-IP and sequencing of RNAs bound by FLAG-tagged RBP in the presence of WT or RNA-binding domain (RBD) mutant of the partner. **(B)** Bar graphs showing the mapped read distribution percentage of the indicated RNA classes in WT, KhpA-3xFLAG, and KhpB-3xFLAG co-IP libraries in the presence of functional and RNA-binding mutants of the partner. Each FLAG-tagged strain is compared to its respective untagged strain. **(C)** Overlap between the RNA targetome of KhpA-3xFLAG (left) and KhpB-3xFLAG (right) in the presence of functional (WT) versus RBD mutant of *khpB* or *khpA*, respectively. Numbers indicate RNAs with a log_2_ fold change ≥ 1, *p*-adj ≤ 0.05, and an average CPM ≥10 in 3xFLAG libraries. **(D)** Comparison of the RNA targetome of KhpA-3xFLAG and KhpB-3xFLAG in the presence of RBD mutants of *khpB* and *khpA*, respectively. Criteria for the numbers indicated are the same as in (C). **(E)** (Top) Violin plot depicting the gene enrichment ratio of KhpA-3xFLAG in the presence of functional (WT) versus R3H-truncated *khpB.* (Bottom) Violin plot depicting the gene enrichment ratio of KhpB-3xFLAG in the presence of functional (WT) versus GXXG-mutated *khpA*. Unique and commonly enriched RNAs are represented by yellow and gray points, respectively. The enrichment ratio for each significantly enriched gene is calculated by dividing the CPM of the tagged library by that of the untagged library.

DESeq2 analysis in the WT partner background identified 323 and 268 significantly enriched transcripts by KhpA and KhpB, respectively, consistent with broad RNA-binding capacity when paired with a functional partner KH-protein (Fig. 5C; Fig. S15B). However, in the partner RBD mutant backgrounds (c*khpB*_ΔR3H and c*khpA*-KH^GDDG^ for KhpA and KhpB, respectively), the enriched RNAs decreased to 148 for KhpA and 92 for KhpB (∼55% and ∼66% reductions; Fig. 5C; Table S4), indicating that RNA binding by one partner strongly relies on the RBD of the other. While KhpA and KhpB shared many targets under WT conditions, this overlap was largely lost with the partner RBD mutant, underscoring the partner’s RNA-binding capacity in sharing specific targets (Fig. 5D; Fig. S15B). Notably, KhpB lost most of both its shared and distinct targets, whereas KhpA showed reduced enrichment for shared targets but retained many KhpA-specific targets (Fig. S15B-C). Global enrichment ratios confirmed widespread reduction in target enrichment in the partner RBD mutant background, with a stronger effect observed for KhpB (Fig. 5E). An interdependency in the RNA targetome enrichment by the two KH-proteins is further supported by hierarchical clustering of log2 fold-changes between different epitope-tagged and untagged KhpA and KhpB pull-downs (Fig. S16). The resulting heatmap revealed co-clustering of KhpA-3xFLAG and KhpB-3xFLAG pull-downs in the WT background, whereas conditions with impaired RNA binding (RBD mutant, Δ partner, and partner RBD mutant backgrounds) formed distinct clusters. Overall, this indicates a shared loss of RNA association when the RBD of either protein is mutated or when the partner protein is deleted.

### RNA-binding domains of KhpA/B and the Jag-N domain of KhpB are crucial for *Campylobacter* adhesion protein target regulation

We next sought to determine the impact of domain mutations on KhpA/B target interactions. Since *capA* mRNA is the most abundantly enriched top target of both KhpA and KhpB (Fig. S4C-D), we decided to use *capA* mRNA as a model mRNA to investigate the functional consequences of the domain mutants. The CapA autotransporter lipoprotein comprises an extracellular passenger domain at the N-terminus and a membrane-anchored C-terminal β-barrel domain. We previously observed an upregulation of CapA protein levels in both Δ*khpA* and Δ*khpB* strains, without significant changes in *capA* mRNA levels (Sharma Lab, unpublished data). Our above listed RIP-seq revealed a loss of *capA* mRNA enrichment in KhpA and KhpB RBD mutants (Fig. 3G) and KhpA in *khpB*_ΔR3H and KhpB in *khpA*_KH^GDDG^ backgrounds (Fig. 6A). To assess effects on *capA* expression, we generated strains expressing CapA-3xFLAG from its native locus in WT and in domain variant strains of 3xFLAG-tagged *khpA* or HA-tagged *khpB*. CapA protein levels, specifically the membrane-anchored β-barrel domain, were upregulated in KhpA_KH^GDDG^, phenocopying Δ*khpA*, whereas KhpA_WT showed WT-similar CapA levels (Fig. 6B, left). While KhpB_KH^GDDA^ and KhpB_WT maintained WT-like CapA levels, consistent with their binding to the *capA* mRNA in these strains, CapA levels increased in the RNA-binding-deficient KhpB_ΔR3H, phenocopying Δ*khpB* (Fig. 6B, right). This suggests that *capA* mRNA binding by both KhpA and KhpB is required for regulating CapA levels. Truncation of the Jag-N domain in KhpB also led to CapA upregulation (Fig. 6B, right), despite retaining *capA* mRNA interaction, indicating that the Jag-N domain of KhpB is also important for *capA* regulation. Together, our data imply that the KH and R3H RNA-binding regions within KhpA and KhpB, respectively, along with the KhpB Jag-N domain, are necessary for *capA* regulation.

**Fig. 6:**
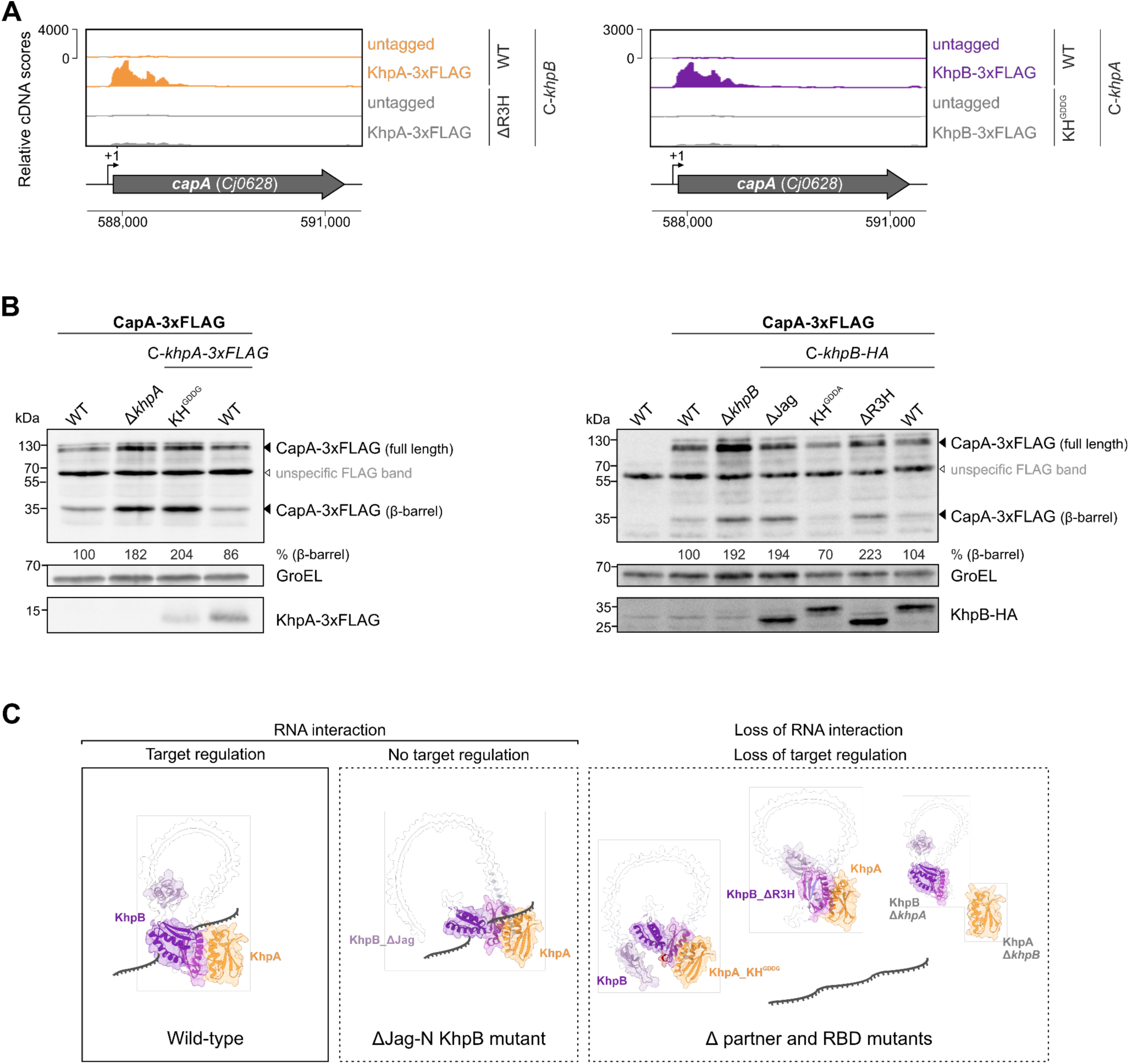
*Campylobacter* adhesion protein expression analyses in the KhpA and KhpB domain mutant backgrounds. **(A)** (Left) Genome browser view depicting cDNA read coverage for KhpA-3xFLAG co-IP libraries in the presence of functional (WT) and R3H-truncated *khpB*, mapped to the *Campylobacter* adhesion protein encoding mRNA (*capA*). (Right) Genome browser view depicting cDNA read coverage for KhpB-3xFLAG co-IP libraries in the presence of functional (WT) and GDDG-mutated *khpA* mapped to *capA*. Each FLAG-tagged strain is compared to its respective untagged strain. **(B)** (Left) Western blot analysis of CapA-3xFLAG expression levels in *C. jejuni* WT, deletion mutants, complemented functional (WT), and domain mutant variants of KhpA-3xFLAG. KhpA-3xFLAG and CapA-3xFLAG were both simultaneously detected with an anti-FLAG antibody. Bands were quantified using ImageJ, normalized to GroEL, and percentages relative to CapA-3xFLAG (β-barrel) in *C. jejuni* WT are given. Blots are indicative of at least three independent replicates. (Right) Western blot analysis of CapA-3xFLAG expression levels in *C. jejuni* WT, deletion mutants, complemented functional (WT), and domain mutant variants of KhpB-HA. CapA-3xFLAG and KhpB-HA were detected with an anti-FLAG and anti-HA antibody, respectively. **(C)** Model summarizing the interplay between *C. jejuni* KhpA and KhpB. In the wild type condition, KhpA and KhpB interact directly and mediate various cellular RNA interactions. Mutation of the KH motif in KhpA and deletion of the non-canonical R3H domain in KhpB disrupts most RNA interactions, suggesting that both conserved and degenerate RBDs of KhpA and KhpB are essential for RNA binding. The RNA interactions of KhpA and KhpB are interdependent: deletion of the partner KH-protein, or mutation of the partner RBD compromises the RNA targetome of KhpA/B. Deletion of either KH-protein or their RBD mutations abolishes *capA* mRNA binding and regulation. In contrast, deletion of the Jag-N domain disrupts *capA* regulation, despite retaining *capA* binding.

## Discussion

In this study, we elucidated the RNA-binding interplay between KhpA and KhpB in *Campylobacter jejuni*, two KH domain proteins that form a heterocomplex. Utilizing transcriptome-wide RIP-seq analyses of WT and mutant strains, we demonstrate that KhpA and KhpB possess overlapping RNA-binding signatures and require each other’s presence to mediate robust interactions with their RNA targets. We also define RNA-binding regions of KhpA and KhpB, highlighting the critical roles of both conserved and degenerate domains in mediating RNA interactions and target regulation (Fig. 6C).

We uncovered that the conserved GXXG motif of KhpA and the R3H domain of KhpB are indispensable for their RNA binding. Despite lacking the canonical RXXXH motif, KhpB seems to primarily rely on its structurally conserved R3H domain for RNA interaction. While conservation analysis of KhpA and KhpB RBD motifs revealed that KhpB invariably lacks an RXXXH motif in the R3H domain across several *Campylobacter* species (Fig. S17), some KhpB homologs harbor a potential functional GKEG motif, instead of the *C. jejuni* GKEA motif. Interestingly, in these orthologues, the asparagine at position 50 (N50) in the GKNG motif of KhpA, which interacts with RNA, is replaced by a negatively charged aspartic acid (D50) (GKDG), likely repelling RNA. Indeed, bacterial three hybrid analysis of *C jejuni* KhpA’s interaction with RNA has recently demonstrated that replacing this asparagine residue with a negatively charged glutamate is sufficient to reduce RNA binding (28). Hence, this amino acid co-variation in the KH motifs of KhpA and KhpB further supports the idea that both RBPs may have co-evolved to fine-tune RNA interactions as a heterocomplex. Recently, crystal structures of RNA-free and RNA-bound *Thermus thermophilus* KhpB (ttKhpB) dimers were reported (29). This revealed that two protomers dimerize via their KH domains to form an RNA-binding groove composed of one R3H and two KH domains, which cooperatively engage a C-rich 10-mer RNA primarily through phosphate backbone contacts. While these *in-vitro* analyses also identified the RXXXH motif in the R3H domain as essential for RNA binding, they excluded KhpA and relied on synthetic RNAs. In contrast, our *in-vivo* data demonstrate that even in the absence of the canonical RXXXH motif, the R3H domain remains central to cognate RNA target recognition.

Our data further suggest that the mutated GKEA motif of KhpB does not directly engage in RNA interaction. While divergent KH domains with altered GXXG motifs have been studied in eukaryotic systems (30), *C. jejuni* KhpB addresses such features in prokaryotic KH proteins. Mutating the conserved glycine residues in the GXXG motif of KH domain proteins can disrupt both RNA binding and structural integrity (31–33). Although alanine in the *C. jejuni* KhpB GKEA motif is chemically similar to glycine, its additional methyl group may introduce steric hindrance and disrupt backbone conformation. We speculate that this alters or reduces the RNA-binding capacity of this motif. Furthermore, as the KH domain of KhpB is likely to interact with KhpA, it is also tempting to speculate that this domain may function as a protein interaction hub, rather than mediating RNA interactions directly (34–36). For KhpA, on the other hand, we observed reduced protein levels upon mutating its GXXG motif, which is consistent with the previously reported reduced protein levels and stability of ProQ and Hfq RNA-binding mutants (37, 38). Given the observed co-variation of GXXG motifs between KhpA and KhpB across *Campylobacter* species, coevolution of the KhpA/B heterocomplex might have optimized their joint RNA-binding activity. KH domain-containing proteins in eukaryotes also typically bind single-stranded RNA or DNA in complex with other RBPs (20). Future studies on high-resolution structures of the KhpA-KhpB complex with RNA would help clarify which residues, both within and beyond the GXXG motifs, make direct RNA contacts. Recent studies in eukaryotic RBPs have also highlighted the RNA-binding potential of IDRs, especially via their extensive positively charged surface (34, 39) and their role in mediating multivalent interactions (40, 41). Thus, the linker IDR region in KhpB, a highly variable region of KhpB across species connecting the Jag-N and KH domains (14) may also contribute to RNA and protein interaction dynamics.

In addition to its KH and R3H domains, KhpB also features a largely uncharacterized N-terminal Jag domain, named after the initial KhpB name (Jag; *spoIII<u>J</u>*-<u>a</u>ssociated <u>g</u>ene) (42). We found Jag-N to be critical for the observed KhpA/B effect on CapA protein expression levels. Although *capA* RNA target enrichment remains intact, truncation of this domain disrupts regulation of the *capA* mRNA target. Also, *in vitro*, KhpB lacking the N-terminal Jag domain together with KhpA was shown to be able to directly bind sRNAs of *Fusobacterium nucleatum* (12). Recent high-throughput screens in yeast and human RBPs have revealed that while RBDs determine RNA-binding and specificity, distinct regulatory domains modulate the expression of target transcripts, primarily through interactions with other proteins (43, 44). The Jag-N domain is likely such a regulatory domain of KhpB that mediates effector contacts essential for efficient target regulation. Although the effectors responsible for this are currently unclear, future work identifying them may shed light on the mode of transcript regulation by KhpA/B.

The impact of RNA interactions on the *in*-*vivo* complex formation of KhpA/B was previously unknown. Using multiple RNA-binding mutants, we demonstrate that *C. jejuni* KhpA interaction with KhpB does not require RNA *in vivo*. This interaction is stable and resistant to RNase treatment, to Jag-N or R3H domain deletions in KhpB, and to point mutations in the GXXG motifs of both KhpA and KhpB. In *S. pneumoniae*, KhpA and KhpB homologs form a complex *in vivo* (*9*). Moreover, a bacterial two-hybrid assay (BACTH) in a heterologous *E. coli* host showed the dimerization ability of *Streptococcus* KhpB’s KH domain with KhpA, as well as a GDDG-mutated KhpA with KhpB (16). The stable KhpA-KhpB complex likely serves multiple functions. First, KhpB impacts KhpA levels, as indicated by the reduced KhpA protein levels in the absence of KhpB. Second, the two proteins rely on each other to maintain their RNA-binding capability. This RNA-binding interdependency is observed at two levels: the presence of the partner protein and an intact RBD of the partner protein.

Such synergy in RNA-binding between two proteins has been reported in eukaryotic proteins (45–48). Our study highlights such interdependence between two prokaryotic RNA-binding proteins. We still note that although GDDG-mutated KhpA and R3H-deleted KhpB barely retained any interaction with RNAs, in this background, the partner KH protein still co-purified a few RNAs. However, the overall lower enrichment ratios in these contexts suggest that the intact KH protein may retain RNA-binding activity, possibly through weak contacts with RNA. The KhpA-KhpB interaction might enhance both affinity and specificity toward most RNA ligands, with KhpA primarily conferring specificity and KhpB, likely through its Jag-N domain, facilitating effector recruitment.

KhpA and KhpB homologs have been previously reported to interact with various RNA classes, including sRNAs, tRNAs, and mRNAs. This led to the hypothesis that KhpA/B may function as sRNA matchmakers in organisms lacking canonical RNA chaperones such as Hfq and ProQ (14). In *C. jejuni,* our RIP-seq data from both wildtype and mutants of KhpA and KhpB point to a primary role for *C. jejuni* KhpA/B in interacting with coding mRNAs. While most of these mRNA interactions depend on the presence of both KH-proteins, a subset of mRNAs still associates with KhpA or KhpB in the absence of their partner. These residual interactions of KhpA and KhpB in the absence of their partner suggest that each protein may retain limited, partner-independent RNA-binding capacity. This could result from participation in distinct ribonucleoprotein (RNP) complexes or from the formation of homooligomers under certain conditions. Supporting this, BACTH assays have shown that KhpA homologs across species can form homomers (16, 28). The stoichiometric balance between KhpA/B hetero- and potential homooligomeric states may vary in response to physiological or environmental cues. Understanding how these complexes shift under different conditions and their impact on target regulation represents an intriguing avenue for future research.

Taken together, our findings demonstrate KhpA and KhpB as interdependent regulators of RNA targets in *C. jejuni*. By revealing their joint RNA-binding determinants, with emphasis on conserved and degenerate RBDs, we expand our understanding of co-regulating RBPs beyond eukaryotic systems. Future research on exploring broader physiological implications and molecular mechanisms of their combinatorial RNA regulation will provide further insights into their functional roles in bacteria.

## Materials and Methods

### Bacterial strains and growth conditions

Strains, plasmids, and DNA oligonucleotides used in this study are listed in Tables S5-S7. *C. jejuni*strains were routinely grown at 37 °C under microaerobic conditions (10% CO_2_, 5% O_2_, 85% N_2_) using a HERAcell 150i incubator (Thermo Fisher Scientific). Cultures were maintained either on Mueller-Hinton (MH) agar or with shaking in Brucella broth (BB), both supplemented with 10 µg/ml vancomycin. Marker-selective antibiotics for solid media included 50 μg/ml kanamycin (Kan), 20 μg/ml chloramphenicol (Cm), 20 μg/ml gentamicin (Gm), or 250 μg/ml hygromycin B (Hyg) when needed. For liquid cultures, BB was inoculated to an initial OD_600_ of 0.005 and incubated at 150 rpm until mid-log phase (OD_600_ 0.4-0.5). Cultures were optionally back-diluted to an OD_600_ of 0.03-0.05 in fresh BB for extended growth. *E. coli* strains were cultured aerobically at 37 °C in Luria-Bertani (LB) broth or on LB agar, with antibiotics added as needed for plasmid maintenance.

### Construction of *C. jejuni* mutant strains

All mutant strains of *C. jejuni* (gene deletions, chromosomal 3xFLAG-tag or HA-tag insertions, and domain mutant constructs expressed from the corresponding native gene locus) were generated through double-crossover homologous recombination, using DNA fragments introduced via electroporation in a *C. jejuni* NCTC11168 background. For selection, antibiotic resistance markers used for cloning included *aac*(*3*)*-IV* (Gm^R^) (49), *aphA-3* (Kan^R^) (50), *C. coli cat* (Cm^R^) (51), or *aph(7”)* (Hyg^R^) (52). All inserted mutations were first validated by colony PCR using Taq polymerase (NEB) and then confirmed by Sanger sequencing (Microsynth). For cloning, Phusion (Thermo Fisher Scientific) or B7 (Biozym) high-fidelity DNA polymerase was used.

### Cloning of deletion mutant strains

Targeted gene deletions of *khpA*/*B* were carried out using a double-crossover homologous recombination approach (53). This method involved replacing most of the coding sequence of the target gene with a non-polar antibiotic resistance cassette flanked by approximately 500 bp of homologous sequence on either side. As an example, deletion of *khpA* (Cj0711) with a non-polar kanamycin (Kan^R^) cassette in a *khpB*-3xFLAG background is described in detail. Approximately 500 bp regions upstream and downstream of *khpA* were amplified from *C. jejuni* NCTC11168 genomic DNA using primers CSO-4317/5222 and CSO-5223/4315, respectively. The reverse primer for the upstream region and the forward primer for the downstream region were designed with 5’ overhangs complementary to the ends of the resistance cassette to allow fusion by overlap extension PCR. The Kan^R^ cassette was amplified from pGG1 (53) using primers HPK1/HPK2. Next, a three-fragment overlap PCR was performed using CSO-4317/4315 on an equimolar ratio of *khpA* upstream, *khpA* downstream, and Kan^R^ cassette. The obtained linear PCR product was electroporated into *C. jejuni* NCTC11168 *khpB*-3xFLAG (CSS-6058) recipient strain as described above. The deletion strain (*khpB*-3xFLAG Δ*khpA*, CSS-7831) was verified using primers CSO-4316/HPK2. For deletions with non-polar Cm^R^, the cassette was amplified using CSO-0613/0614 from pGD107.1 (53).

### Construction of C-terminally epitope-tagged strains

C-terminal epitope tagging was performed by integrating a 3xFLAG-or HA-tag at the native locus using homologous recombination. The tagging strategy involved overlap PCR and is briefly explained for the *C. jejuni* NCTC11168 KhpB-HA strain (CSS-5898). Approximately 500 bp upstream of *khpB* penultimate codon was amplified using primers CSO-4304/4352, and 500 bp downstream of *khpB* was amplified with HPK2/CSO-4307. An HA tag fused to a non-polar Gm^R^ cassette was amplified from pUC1813-apra (49) using CSO-3560/HPK2. All three fragments (upstream, cassette, and downstream) were mixed in equimolar amounts and assembled by overlap PCR using primers CSO-4304/4307. Subsequent clones were validated by colony PCR (CSO-4308/HPK2) and Sanger sequencing (CSO-6293).

### Cloning of KhpA and KhpB domain mutant variants

For domain mutation of *khpA* and *khpB*, a copy of the respective domain mutant variants was introduced into the native locus of the corresponding deletion strains (Δ*khpA* and Δ*khpB*, respectively) to allow homologous recombination. Domain mutant constructs were first created in plasmids by subcloning the *C. jejuni* sequence into two consecutive plasmids. First, a plasmid vector backbone expressing *khpA* or *khpB* was created by integrating full-length gene constructs into commercially available pGEM®-T Easy Vector Systems (Promega, #A1360). Second, site-directed mutagenesis was carried out on these vector backbones using the Q5 site-directed mutagenesis kit (NEB #E0554S) to introduce different domain mutations. As an example, the construction of C-terminally 3xFLAG-tagged KhpA_KH^GDDG^ mutant is described in detail. First, the ∼500 bp upstream and downstream of *khpA* were amplified using primers CSO-4317/HPK1 and CSO-5223/4315, respectively, on the published *khpA*::3xFLAG (CSS-6060) strain (Sharma Lab, unpublished). The non-polar Gm^R^ cassette was amplified from pUC1813-apra (49) using primers HPK1/HPK2. The three fragments with 3xFLAG overhangs were mixed together in equimolar ratios and amplified by overlap PCR using primers CSO-4317/4315. After confirming the correct size by gel electrophoresis, the linear PCR product was 3’ A-tailed using Taq polymerase (NEB) and ligated into pGEM®-T Easy Vector following the manufacturer’s instructions. The ligation product was transformed into competent *E. coli* Top10 cells. Clones/plasmids of the resulting pMN1 were verified by colony PCR (CSO-4317/HPK2) and Sanger sequencing (CSO-4317/6293). Second, pMN1.2 was used as a template to introduce point mutations corresponding to KhpA_KH^GDDG^ by Q5 site-directed mutagenesis using non-overlapping primers CSO-6739/6740 according to the manufacturer’s instructions (NEB #E0554S). This resulted in plasmid pMN7.1. Clones/plasmids were verified by colony PCR (CSO-4317/HPK2) and Sanger sequencing (CSO-6293). The construct containing KhpA_KH^GDDG^::3xFLAG with a non-polar Gm^R^ cassette was amplified from the subsequent plasmid pMN7.1 using CSO-4317/4315, and the linear PCR product was electroporated into Δ*khpA* (CSS-7299). Recovered clones were verified by colony PCR (CSO-4316/HPK2) and Sanger sequencing (CSO-6293).

A similar strategy was employed to generate untagged versions of the respective domain mutants with a few changes. As an example, the construction of an untagged KhpA_KH^GDDG^ mutant is described in detail. To remove the C-terminal 3xFLAG-tag, inverse PCR was performed on pMN1.2 using non-overlapping primers CSO-6591/6594. Purified DNA was *Dpn*I digested, self-ligated, and transformed into *E. coli* Top10. The resulting plasmid pMN5.1 was sequenced with CSO-6293 and used as a template for Q5 site-directed mutagenesis using CSO-6739/6740 as described above. The subsequent plasmid pMN8.6 was sequenced with CSO-6293 and used as a template to introduce KhpA_KH^GDDG^ with a non-polar Gm^R^ cassette into *C. jejuni* Δ*khpA* (CSS-7299).

### Transformation of *C. jejuni* by electroporation

All *C. jejuni* mutants used in this study were generated by electroporation of DNA fragments into recipient strains. Strains were streaked out on MH agar plates until passage one and resuspended in cold electroporation solution (272 mM sucrose, 15% (w/v) glycerol). The bacterial suspension was spun down (6,500x*g*, 5 min, 4 °C), washed twice with the same solution, and resuspended based on pellet size (100-300 μl). For each transformation, 50 μl of the cell suspension was mixed with 500-1000 ng of purified PCR product. Cells were electroporated in a 1 mm gap cuvette (Cell Projects) at 2.5 kV for 4-6 ms using a Biorad MicroPulser™. Cells were recovered in 100 μl BB medium and plated on a non-selective MH-agar plate containing vancomycin and incubated overnight at 37 °C under microaerobic conditions. The next day, cells were restreaked on a marker-selective MH agar plate and incubated until colonies appeared (2-5 days). Clones were verified by colony PCR and, where applicable, by Sanger sequencing.

### RNA immunoprecipitation

Co-IP combined with RNA-seq (RIP-seq) with different FLAG-tagged strains was performed as previously described for *C. jejuni* NCTC11186 (54). Strains expressing 3xFLAG tagged WT or mutant versions of KhpA or KhpB, along with respective untagged controls, were cultured in 50 ml BB containing 10 µg/ml vancomycin until mid-exponential phase (OD_600_ ≈ 0.6). Cells were harvested by centrifugation at 4,000x*g* for 20 min at 4 °C, resuspended in 1 ml of Buffer A (20 mM Tris-HCl, pH 8.0, 150 mM KCl, 1 mM MgCl₂, 1 mM DTT), and centrifuged again (11,000g, 3 min, 4 °C). In parallel, 1 ml of the culture was harvested for whole cell lysate analysis and resuspended in 1x protein loading buffer (62.5 mM Tris-HCl, pH 6.8, 100 mM DTT, 10% (v/v) glycerol, 2% (w/v) SDS, 0.01% (w/v) bromophenol blue) and boiled at 95 °C for 8 min with shaking at 1,000 rpm. Cell pellets were snap-frozen in liquid nitrogen and stored at -80 °C. Cell pellets were thawed on ice and resuspended in 1 ml of lysis buffer (957 μl Buffer A, 1 μl 1 mM DTT, 10 μl 0.1 M PMSF, 2 μl Triton X-100, 20 μl DNase I, and 10 μl Superase-In RNase inhibitor). The suspension was transferred to FastPrep tubes and subjected to mechanical disruption using a FastPrep homogenizer (MP-Biomedicals, matrix B) at 4 M/s for 15 seconds. Lysates were then clarified by centrifugation at 13,000 rpm for 10 min at 4 °C, and an aliquot (1 OD_600_) was collected as input/lysate control for subsequent western blot analysis. Clarified lysate was then incubated with 35 μl of anti-FLAG antibody (monoclonal anti-FLAG M2, Sigma, #F1804) for 30 min at 4 °C on a rocker. Protein A-Sepharose beads (75 μl/sample) (Sigma, #P6649), pre-washed and equilibrated in Buffer A, were then added and incubated for an additional 30 min with gentle rocking at 4 °C. Beads were washed five times (15,000x*g*, 1 min, 4 °C) with 500 μl Buffer A to remove non-specifically bound material. Finally, 500 μl Buffer A was added to the beads. Bound RNA and protein were extracted using 500 μl of phenol-chloroform-isoamyl alcohol (P:C:I), followed by 5 min incubation at room temperature (RT) and centrifugation at 13,000 rpm for 30 min at 4 °C. To recover RNA, the aqueous phase was precipitated overnight using a 30:1 mix (ethanol:3M sodium acetate) plus 1 μl of GlycoBlue at -20 °C. To recover protein, beads in the organic phase were treated with 1.4 ml ice-cold acetone and precipitated overnight at -20 °C. The next day, RNA samples were centrifuged at 13,000 rpm for 30 min at 4 °C and washed with 500 μl of ice-cold ethanol. After removing residual ethanol, the pellet was air dried and resuspended in 30 μl H_2_O (5 min, 65 °C). Protein samples were centrifuged at 15,000 rpm for 30 min at 4 °C, washed twice with acetone, and resuspended in 120 μl of 1x protein loading buffer (95 °C, 8 min). Eluted proteins and RNA were used for downstream western blot or RNA-seq analysis, respectively.

### cDNA library preparation

RIP RNA samples were treated with DNase I (ThermoFisher Scientific) to remove residual gDNA. cDNA library preparation used an oligonucleotide adapter which was ligated to the 3’ end of the RNA molecules. First-strand cDNA synthesis was performed using M-MLV reverse transcriptase with the 3’ adapter as primer. After purification, the 5’ Illumina TruSeq sequencing adapter was ligated to the 3’ end of the antisense cDNA. The resulting cDNA was PCR-amplified using a high-fidelity DNA polymerase, and the barcoded TruSeq libraries were pooled in approximately equimolar amounts. Sequencing of pooled libraries, spiked with PhiX control library, was performed at a minimum of 4 million reads per sample in single-end mode with 100 cycles on the NextSeq 2000 platform (Illumina). Demultiplexed FASTQ files were generated with bcl-convert v4.2.4 or 4.3.6 (Illumina).

### RIP-seq data analysis

Raw sequencing reads were subjected to quality and adapter trimming via Cutadapt (55) v2.5 using a cutoff Phred score of 20 and discarding reads without any remaining bases (parameters: --nextseq-trim=20 -m 1 -a AGATCGGAAGAGCACACGTCTGAACTCCAGTCAC). Afterwards, all reads longer than 11 nt were aligned to the *C. jejuni* NCTC 11168 reference genome (RefSeq assembly accession: GCF_000009085.1) using the pipeline READemption (56) v2.0.3 with segemehl v0.3.4 (57) and an accuracy cut-off of 95% (parameters: -l 12 -a 95). READemption gene_quanti was applied to quantify aligned reads overlapping genomic features by at least 10 nt on the sense strand based on NCBI gene annotations complemented with previously determined 5′ UTRs and small RNAs (58) (parameters: --min_overlap 10 --features “5’-UTR, CDS, rRNA, tRNA, transcript,sRNA“). Based on these counts, differential expression analysis was conducted via DESeq2 (59) v1.24.0. Read counts were normalized by DESeq2, and fold-change shrinkage was conducted by setting the parameter betaPrior to TRUE. Enrichment of tagged vs. control libraries was tested using parameter altHypothesis=“greater” for the results function. Significant features were selected based on thresholds for adjusted *p*-value after Benjamini-Hochberg correction (*p*-adj) ≤0.05 and log2FoldChange ≥1. An additional enrichment cutoff feature of average counts per million ≥ 10 in both replicates of the 3xFLAG tagged libraries was set based on normalization constants derived from the total number of reads mapped to the respective libraries. Integrated genome browser (IGB) (60) was used to visualize the cDNA read coverages.

### Protein gel electrophoresis and immunoblotting

*C. jejuni* cells were cultured to mid-exponential phase (OD_600_ = 0.4-0.5), unless a different OD_600_ is specified, and harvested by centrifugation at 13,000 rpm for 3 min. Cell pellets were resuspended in 1x protein loading buffer (62.5 mM Tris-HCl, pH 6.8, 100 mM DTT, 10% (v/v) glycerol, 2% (w/v) SDS, 0.01% (w/v) bromophenol blue) and boiled at 95 °C for 8 min with shaking at 1,000 rpm. Samples equivalent to an OD_600_ of 0.1 for whole cell lysates, or 10 in case of co-IP eluate samples, were separated on 10 % or 12 % SDS-polyacrylamide (PAA) gels and transferred to nitrocellulose membranes by semi-dry blotting (Peqlab; 90 min at 0.34 A/blot). In some cases, to better resolve lower molecular weight protein bands, NuPAGE Bis-Tris gel systems (Thermo Fisher Scientific, #NP0321) were used according to the manufacturer’s instructions, and samples equivalent to an OD_600_ of 0.05 were loaded. Membranes were blocked for 1 h with 10 % (w/v) milk powder in TBS-T (Tris-buffered saline-Tween-20), then incubated overnight at 4 °C with primary antibody in 3 % BSA/TBS-T [monoclonal mouse anti-FLAG (1:1,000; Sigma-Aldrich, #F1804-1MG); or monoclonal mouse anti-HA (1:1,000; Biolegend, #B901513)]. After washing (TBS-T), membranes were incubated for 1 h at room temperature with HRP-conjugated secondary antibody [sheep polyclonal, anti-mouse IgG horseradish peroxidase (HRP) conjugate (1:10,000 GE Healthcare, #RPN4201)]. Blots were developed using ECL-reagent (luminol, para-hydroxycomaric acid, and H_2_O_2_ in Tris-HCl buffer) and imaged with an ImageQuant LAS-4000 Imager (GE Healthcare, version 1.3, build 1.3.0.134). For loading control, membranes were probed with polyclonal rabbit anti-GroEL (1:10,000; Sigma-Aldrich, #G6532-5ML) and HRP-conjugated goat polyclonal anti-rabbit IgG (1:10,000; GE Healthcare, #RPN4301). Prestained protein marker (Thermo Fisher Scientific) was used for molecular weight estimation. Bands were quantified using ImageJ (61).

### Protein-protein co-IP

Co-IP to investigate protein-protein interactions was performed as described for the RIP-seq protocol (see above) with a few modifications. DYKDDDDK Fab-Trap agarose beads (Chromtek) were used in place of anti-FLAG antibody and Protein A-Sepharose beads, with bead equilibration and washing carried out according to the manufacturer’s instructions. After the last washing step, the beads were resuspended in 60 μl of 1x protein loading buffer and boiled at 95 °C for 8 min with shaking at 1,000 rpm. For RNase A-treated co-IPs, no Superase-In RNase inhibitor was used in the lysis buffer, and 2 μl of RNase A (10 mg/ml; Thermo Fischer Scientific #EN0531) was added to the lysates of treated samples and incubated for 30 minutes at 4° C with gentle rocking, after which co-IP was performed. Lysate samples and eluate samples from all co-IP experiments were used for subsequent western blot analysis.

### Protein complex and structure prediction

The wild-type or domain variants of KhpA and KhpB were modelled together as a complex using ColabFold software (62) based on AlphaFold2 (24) and MMseqs2 (63) with default parameters. Models with the best ranking and high ipTM scores were selected. Structural predictions were visualized using UCSF ChimeraX (64).

### pyRBDome pipeline

Protein sequences corresponding to KhpA and KhpB were retrieved from UniProt for the following bacterial species: *Campylobacter jejuni* NCTC11168 (KhpA: Q0PAG8; KhpB: Q0P9U2), *Streptococcus pneumoniae* serotype 2 strain D39/NCTC 7466 (KhpA: A0A0H2ZMB4; KhpB: A0A0H2ZPS7), *Clostridioides difficile* strain 630 (KhpA: Q18BB7; KhpB: Q181S9), *Helicobacter pylori* strain G27 (KhpA: B5Z8E6; KhpB: B5Z968). For the pyRBDome analyses, Alphafold models were submitted to the pyRBDome pipeline (26). These were analysed using BindUP (65) to identify positively charged amino acid patches, RNABindRPlus (66) to predict RNA-binding amino acids based on sequence conservation, PST-PRNA (67) to predict RNA-binding amino acids based on surface topology and sequence conservation, DisoRDPBind (68) to predict RNA-binding amino acids in disordered regions, and FTMap (69) to identify small molecule binding pockets. The prediction results were then analysed by an XGBoost ensemble model that had been trained on outputs from these tools on human RBPs to produce a final RNA-binding probability score. Note that bacterial RNA-binding proteins were not included in the training of the XGBoost model.

### Bacterial two-hybrid (B2H) assays

To construct plasmids for B2H analysis, sequences encoding *Cje* KhpA and KhpB were PCR amplified from genomic DNA with primers oSJ1+oSJ2 (KhpA) and oSJ5+oSJ6 (KhpB) and ligated into pPrey and pBait constructs (α-NTD and λCI, respectively; (70) using NotI and BamHI restriction sites. To carry out B2H experiments, *E. coli* reporter cells (KB473; (71)) were transformed with combinations of pPrey and pBait plasmids encoding hybrid constructs with both KhpA and KhpB as Bait and Prey. For negative controls, transformations included one of the hybrid constructs and a corresponding empty construct (α-empty or CI-empty). Transformations and bacterial growth were conducted in 96-well plates as previously described (72). LB medium in overnight cultures was supplemented with 100 μg/ml carbenicillin, chloramphenicol and kanamycin, and back-dilutions contained an additional 5 μM isopropyl β-D-1-thiogalactopyranoside (IPTG). β-galactosidase (LacZ) activity was measured from lysed mid-log cells as previously described (72). Two-hybrid interactions were quantified by calculating the fold-stimulation over basal LacZ levels: i.e. LacZ activity in the presence of both hybrid constructs (e.g. pPrey-KhpA and pBait-KhpB) divided by the highest activity observed in negative control transformations containing one hybrid component and one empty construct. Bar plots represent the mean and standard deviation (SD) of fold-stimulation values calculated across 6 replicates of 3 independent experiments.

### Text editing

The manuscript text was reviewed using Grammarly (https://app.grammarly.com/) to ensure grammatical accuracy.

## Data availability

The raw reads of all RIP-seq libraries have been deposited in the NCBI Gene Expression Omnibus (GEO) under the accession number XXX (link).

## Acknowledgements

We thank members of the Sharma lab for critical discussions on the manuscript. We thank Ann-Kristin Herber for assistance with RNaseA-treated co-IPs and Elena Katzowitzsch from the Core Unit SysMed at the University of Würzburg for technical support. We thank Sungeun Jo and Chandra Gravel for assistance in cloning constructs for B2H analysis.

## Author Contributions

M.N., S.S., and C.M.S. designed the experiments. M.N. and S.S. performed lab work and analyzed the data. T.B. and T.G. performed sequencing and bioinformatic analyses. S.G. performed pyRBDome analyses. N.W.L. and K.B. collected and analyzed two-hybrid data. M.N., M.A., and C.M.S. interpreted the data and wrote the manuscript, with comments from co-authors. C.M.S. supervised research and provided resources.

## References

1. M. Corley, M. C. Burns, G. W. Yeo, How RNA-Binding Proteins Interact with RNA: Molecules and Mechanisms. Mol. Cell 78, 9–29 (2020).

2. S. H. Stitzinger, S. Sohrabi-Jahromi, J. Söding, Cooperativity boosts affinity and specificity of proteins with multiple RNA-binding domains. NAR Genom. Bioinform. 5, lqad057 (2023).

3. C. D. Mackereth, M. Sattler, Dynamics in multi-domain protein recognition of RNA. Curr. Opin. Struct. Biol. 22, 287–296 (2012).

4. B. M. Lunde, C. Moore, G. Varani, RNA-binding proteins: modular design for efficient function. Nat. Rev. Mol. Cell Biol. 8, 479–490 (2007).

5. E. Ng Kwan Lim, C. Sasseville, M.-C. Carrier, E. Massé, Keeping up with RNA-based regulation in bacteria: New roles for RNA binding proteins. Trends Genet. 37, 86–97 (2021).

6. P. Babitzke, Y.-J. Lai, A. J. Renda, T. Romeo, Posttranscription initiation control of gene expression mediated by bacterial RNA-binding proteins. Annu. Rev. Microbiol. 73, 43–67 (2019).

7. E. Holmqvist, J. Vogel, RNA-binding proteins in bacteria. Nat. Rev. Microbiol. 16, 601–615 (2018).

8. A. P. Quendera, et al., RNA-binding proteins driving the regulatory activity of small non-coding RNAs in bacteria. Front. Mol. Biosci. 7, 78 (2020).

9. J. J. Zheng, A. J. Perez, H.-C. T. Tsui, O. Massidda, M. E. Winkler, Absence of the KhpA and KhpB (JAG/EloR) RNA-binding proteins suppresses the requirement for PBP2b by overproduction of FtsA in *Streptococcus pneumoniae* D39. Mol. Microbiol. 106, 793–814 (2017).

10. V. Lamm-Schmidt, et al., Grad-seq identifies KhpB as a global RNA-binding protein in *Clostridioides difficile* that regulates toxin production. microLife 2, uqab004 (2021).

11. C. Michaux, M. Gerovac, E. E. Hansen, L. Barquist, J. Vogel, Grad-seq analysis of Enterococcus faecalis and Enterococcus faecium provides a global view of RNA and protein complexes in these two opportunistic pathogens. microLife 4, uqac027 (2023).

12. Y. Zhu, F. Ponath, V. Cosi, J. Vogel, A global survey of small RNA interactors identifies KhpA and KhpB as major RNA-binding proteins in *Fusobacterium nucleatum*. Nucleic Acids Res. 52, 3950–3970 (2024).

13. R. Han, et al., RNA Binding Proteins KhpA and KhpB Interact with Small Regulatory RNAs and Affect Global Gene Expression in *Deinococcus radiodurans*. BioRxiv (2025) 10.1101/2025.09.18.676964.

14. M. Olejniczak, X. Jiang, M. M. Basczok, G. Storz, KH domain proteins: Another family of bacterial RNA matchmakers? Mol. Microbiol. 117, 10–19 (2022).

15. G. A. Stamsås, et al., Identification of EloR (Spr1851) as a regulator of cell elongation in *Streptococcus pneumoniae*. Mol. Microbiol. 105, 954–967 (2017).

16. A. R. Winther, M. Kjos, G. A. Stamsås, L. S. Håvarstein, D. Straume, Prevention of EloR/KhpA heterodimerization by introduction of site-specific amino acid substitutions renders the essential elongasome protein PBP2b redundant in *Streptococcus pneumoniae*. Sci. Rep. 9, 3681 (2019).

17. A. Tikhomirova, et al., *Campylobacter jejuni* virulence factors: update on emerging issues and trends. J. Biomed. Sci. 31, 45 (2024).

18. P. M. Burnham, D. R. Hendrixson, *Campylobacter jejuni*: collective components promoting a successful enteric lifestyle. Nat. Rev. Microbiol. 16, 551–565 (2018).

19. J. Parkhill, et al., The genome sequence of the food-borne pathogen *Campylobacter jejuni* reveals hypervariable sequences. Nature 403, 665–668 (2000).

20. R. Valverde, L. Edwards, L. Regan, Structure and function of KH domains. FEBS J. 275, 2712– 2726 (2008).

21. N. V. Grishin, KH domain: one motif, two folds. Nucleic Acids Res. 29, 638–643 (2001).

22. J. Pereira, A. N. Lupas, The ancestral KH peptide at the root of a domain family with three different folds. Bioinformatics 34, 3961–3965 (2018).

23. N. V. Grishin, The R3H motif: a domain that binds single-stranded nucleic acids. Trends Biochem. Sci. 23, 329–330 (1998).

24. J. Jumper, et al., Highly accurate protein structure prediction with AlphaFold. Nature 596, 583–589 (2021).

25. J. R. Parrish, et al., A proteome-wide protein interaction map for *Campylobacter jejun*i. Genome Biol. 8, R130 (2007).

26. L.-C. Chu, et al., pyRBDome: a comprehensive computational platform for enhancing RNA-binding proteome data. Life Sci. Alliance 7 (2024).

27. D. Hollingworth, et al., KH domains with impaired nucleic acid binding as a tool for functional analysis. Nucleic Acids Res. 40, 6873–6886 (2012).

28. K. T. Nguyen, et al., Molecular genetic characterization of bacterial KH-domain proteins. BioRxiv (2026) 10.64898/2026.03.16.712068.

29. K. Fukui, T. Murakawa, S. Baba, T. Kumasaka, T. Yano, KH-R3H domain cooperation in RNA recognition by the global RNA-binding protein KhpB. Nat. Commun. 16, 8028 (2025).

30. M. A. Brykailo, A. H. Corbett, J. L. Fridovich-Keil, Functional overlap between conserved and diverged KH domains in *Saccharomyces cerevisiae* SCP160. Nucleic Acids Res. 35, 1108–1118 (2007).

31. J. García-Mena, A. Das, A. Sánchez-Trujillo, C. Portier, C. Montañez, A novel mutation in the KH domain of polynucleotide phosphorylase affects autoregulation and mRNA decay in *Escherichia coli*. Mol. Microbiol. 33, 235–248 (1999).

32. M. Yadav, et al., The KH domain facilitates the substrate specificity and unwinding processivity of DDX43 helicase. J. Biol. Chem. 296, 100085 (2021).

33. A. G. Wong, K. L. McBurney, K. J. Thompson, L. M. Stickney, G. A. Mackie, S1 and KH domains of polynucleotide phosphorylase determine the efficiency of RNA binding and autoregulation. J. Bacteriol. 195, 2021–2031 (2013).

34. F. Bonneau, et al., Nuclear mRNPs are compact particles packaged with a network of proteins promoting RNA-RNA interactions. Genes Dev. 37, 505–517 (2023).

35. S. Fribourg, D. Gatfield, E. Izaurralde, E. Conti, A novel mode of RBD-protein recognition in the Y14-Mago complex. Nat. Struct. Biol. 10, 433–439 (2003).

36. C. L. Kielkopf, N. A. Rodionova, M. R. Green, S. K. Burley, A novel peptide recognition mode revealed by the X-ray structure of a core U2AF35/U2AF65 heterodimer. Cell 106, 595–605 (2001).

37. Y. El Mouali, et al., Scanning mutagenesis of RNA-binding protein ProQ reveals a quality control role for the Lon protease. RNA 27, 1512–1527 (2021).

38. A. Zhang, D. J. Schu, B. C. Tjaden, G. Storz, S. Gottesman, Mutations in interaction surfaces differentially impact E. coli Hfq association with small RNAs and their mRNA targets. J. Mol. Biol. 425, 3678–3697 (2013).

39. I. Zigdon, et al., Beyond RNA-binding domains: determinants of protein-RNA binding. RNA 30, 1620–1633 (2024).

40. M. Macossay-Castillo, et al., The Balancing Act of Intrinsically Disordered Proteins: Enabling Functional Diversity while Minimizing Promiscuity. J. Mol. Biol. 431, 1650–1670 (2019).

41. A. S. Holehouse, B. B. Kragelund, The molecular basis for cellular function of intrinsically disordered protein regions. Nat. Rev. Mol. Cell Biol. 25, 187–211 (2024).

42. J. Errington, et al., Structure and function of the *spoIIIJ* gene of *Bacillus subtilis*: a vegetatively expressed gene that is essential for sigma G activity at an intermediate stage of sporulation. J. Gen. Microbiol. 138, 2609–2618 (1992).

43. K. Reynaud, A. M. McGeachy, D. Noble, Z. A. Meacham, N. T. Ingolia, Surveying the global landscape of post-transcriptional regulators. Nat. Struct. Mol. Biol. 30, 740–752 (2023).

44. A. R. Thurm, et al., High-throughput discovery of regulatory effector domains in human RNA-binding proteins. BioRxiv (2024) 10.1101/2024.07.19.604317.

45. P. Chen, et al., A Csde1-Strap complex regulates plasma cell differentiation by coupling mRNA translation and decay. Nat. Commun. 16, 2906 (2025).

46. G. Behrens, V. Heissmeyer, Cooperation of RNA-Binding Proteins -a Focus on Roquin Function in T Cells. Front. Immunol. 13, 839762 (2022).

47. S. Winterbourne, et al., Integrative structural analysis of NF45-NF90 heterodimers reveals architectural rearrangements and oligomerization on binding dsRNA. Nucleic Acids Res. 53 (2025).

48. L. Voith von Voithenberg, et al., Recognition of the 3’ splice site RNA by the U2AF heterodimer involves a dynamic population shift. Proc Natl Acad Sci USA 113, E7169–E7175 (2016).

49. S. Bury-Moné, et al., Presence of active aliphatic amidases in *Helicobacter* species able to colonize the stomach. Infect. Immun. 71, 5613–5622 (2003).

50. S. Skouloubris, J. M. Thiberge, A. Labigne, H. De Reuse, The *Helicobacter pylori* UreI protein is not involved in urease activity but is essential for bacterial survival *in vivo*. Infect. Immun. 66, 4517–4521 (1998).

51. Y. Wang, D. E. Taylor, Chloramphenicol resistance in *Campylobacter coli*: nucleotide sequence, expression, and cloning vector construction. Gene 94, 23–28 (1990).

52. A. Cameron, E. C. Gaynor, Hygromycin B and apramycin antibiotic resistance cassettes for use in *Campylobacter jejuni*. PLoS ONE 9, e95084 (2014).

53. G. Dugar, et al., The CsrA-FliW network controls polar localization of the dual-function flagellin mRNA in *Campylobacter jejuni*. Nat. Commun. 7, 11667 (2016).

54. S. Sharma, C. M. Sharma, Identification of RNA Binding Partners of CRISPR-Cas Proteins in Prokaryotes Using RIP-Seq. Methods Mol. Biol. 2404 111–133 (2022).

55. M. Martin, Cutadapt removes adapter sequences from high-throughput sequencing reads. EMBnet j. 17, 10 (2011).

56. K. U. Förstner, J. Vogel, C. M. Sharma, READemption-a tool for the computational analysis of deep-sequencing-based transcriptome data. Bioinformatics 30, 3421–3423 (2014).

57. S. Hoffmann, et al., Fast mapping of short sequences with mismatches, insertions and deletions using index structures. PLoS Comput. Biol. 5, e1000502 (2009).

58. G. Dugar, et al., High-resolution transcriptome maps reveal strain-specific regulatory features of multiple *Campylobacter jejuni* isolates. PLoS Genet. 9, e1003495 (2013).

59. M. I. Love, W. Huber, S. Anders, Moderated estimation of fold change and dispersion for RNA-seq data with DESeq2. Genome Biol. 15, 550 (2014).

60. N. H. Freese, D. C. Norris, A. E. Loraine, Integrated genome browser: visual analytics platform for genomics. Bioinformatics 32, 2089–2095 (2016).

61. C. A. Schneider, W. S. Rasband, K. W. Eliceiri, NIH Image to ImageJ: 25 years of image analysis. Nat. Methods 9, 671–675 (2012).

62. M. Mirdita, et al., ColabFold: making protein folding accessible to all. Nat. Methods 19, 679– 682 (2022).

63. M. Steinegger, J. Söding, MMseqs2 enables sensitive protein sequence searching for the analysis of massive data sets. Nat. Biotechnol. 35, 1026–1028 (2017).

64. E. F. Pettersen, et al., UCSF ChimeraX: structure visualization for researchers, educators, and developers. Protein Sci. 30, 70–82 (2021).

65. D. Alexandrovich, S. Kagan, Y. Mandel-Gutfreund, BindUP-Alpha: A Webserver for Predicting DNA-and RNA-binding Proteins based on Experimental and Computational Structural Models☆. J. Mol. Biol. 437, 169240 (2025).

66. R. R. Walia, et al., RNABindRPlus: a predictor that combines machine learning and sequence homology-based methods to improve the reliability of predicted RNA-binding residues in proteins. PLoS ONE 9, e97725 (2014).

67. P. Li, Z.-P. Liu, PST-PRNA: prediction of RNA-binding sites using protein surface topography and deep learning. Bioinformatics 38, 2162–2168 (2022).

68. C. J. Oldfield, Z. Peng, L. Kurgan, Disordered RNA-Binding Region Prediction with DisoRDPbind. Methods Mol. Biol. 2106, 225–239 (2020).

69. D. Kozakov, et al., The FTMap family of web servers for determining and characterizing ligand-binding hot spots of proteins. Nat. Protoc. 10, 733–755 (2015).

70. S. L. Dove, J. K. Joung, A. Hochschild, Activation of prokaryotic transcription through arbitrary protein-protein contacts. Nature 386, 627–630 (1997).

71. K. E. Berry, A. Hochschild, A bacterial three-hybrid assay detects *Escherichia coli* Hfq-sRNA interactions *in vivo*. Nucleic Acids Res. 46, e12 (2018).

72. O. M. Stockert, C. M. Gravel, K. E. Berry, A bacterial three-hybrid assay for forward and reverse genetic analysis of RNA-protein interactions. Nat. Protoc. 17, 941–961 (2022).

