## Supplementary Figures for "Interdependent mode of RNA recognition by the interacting KH-domain proteins KhpA and KhpB"

**This document contains:**

**Fig. S1-S17**

**Supplementary References**

### Supplementary Figures

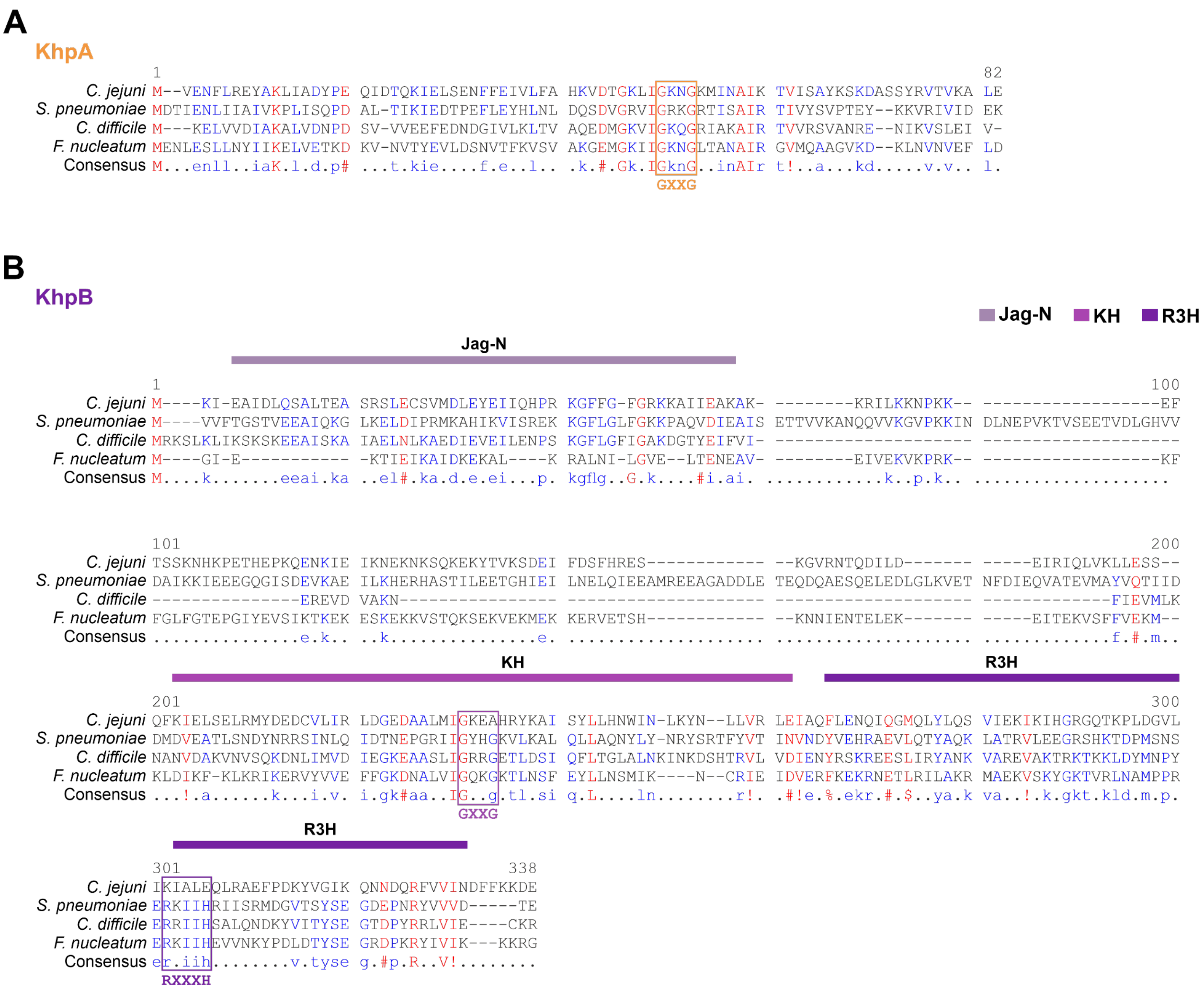


**Fig. S1: Multiple sequence alignment of KhpA and KhpB across different bacteria. (A-B)** Alignment of protein sequences of KhpA (A) and KhpB (B) from *Campylobacter jejuni*, *Streptococcus pneumoniae*, *Clostridium difficile*, *and Fusobacterium nucleatum* using Multalin (1). Different domains and the RNA-binding motifs (GXXG motif of the KH domain and the RXXXH motif of the R3H domain) are highlighted.

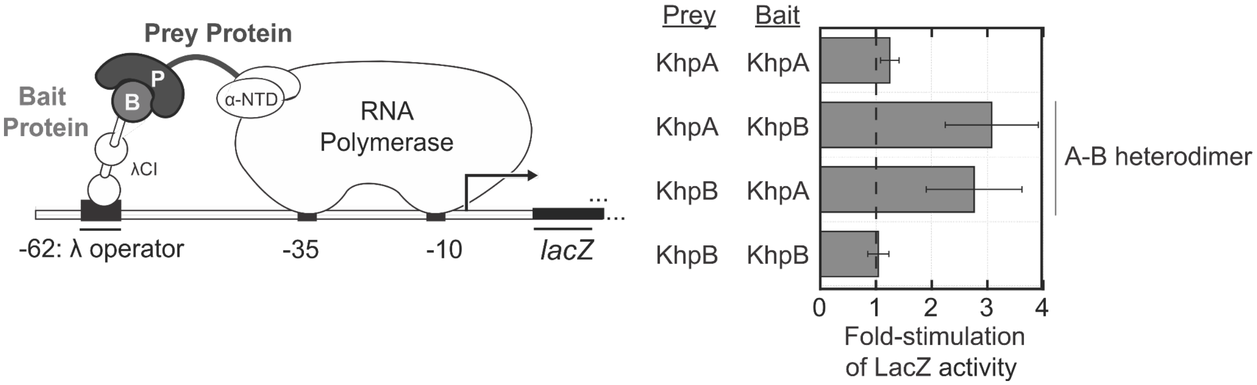


**Fig. S2: Bacterial two-hybrid (B2H) assays with *C. jejuni* KhpA and KhpB in the heterologous *E. coli*.** (Left) Schematic representation of the bacterial two-hybrid (B2H) system (2). Interaction between Prey (P) and Bait (B) proteins, fused respectively to the N-terminal domain of the RNA polymerase α-subunit (α-NTD) and the bacteriophage λ CI protein (λCI), activates transcription of the lacZ reporter gene in reporter strain KB473 (3). (Right) Bar graphs showing the B2H results for potential KhpA and KhpB hetero- and homodimerization. Bars represent fold-stimulation of LacZ (-galactosidase) activity when both Bait and Prey protein are present relative to basal levels measured for the corresponding single-protein negative control (each pPrey or pBait construct transformed alongside an empty construct).

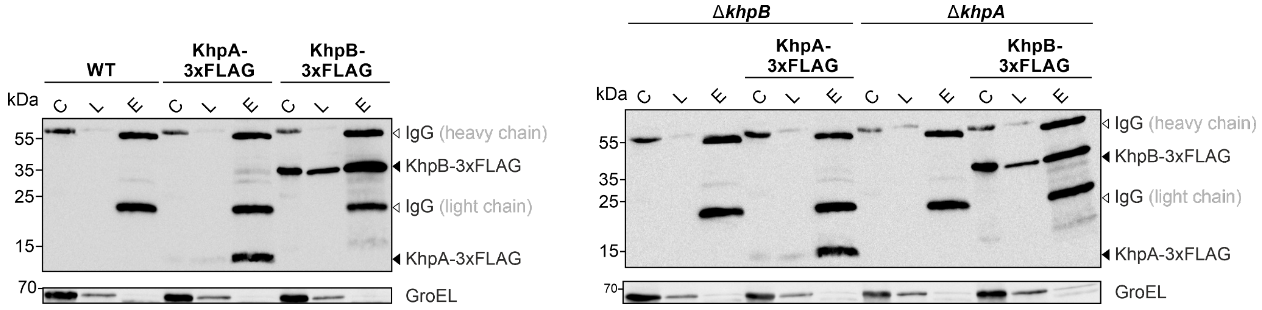


**Fig. S3: Co-IP results of KhpA and KhpB in the presence and absence of partner KH-protein.** Western blot analysis of immunoprecipitated proteins from *C. jejuni* lysates in the WT background (left) and Δ partner background (right). FLAG-tagged proteins were detected using an anti-FLAG antibody. C: culture, L: lysate, E: eluate. GroEL served as a loading control. One representative biological replicate out of two is shown.

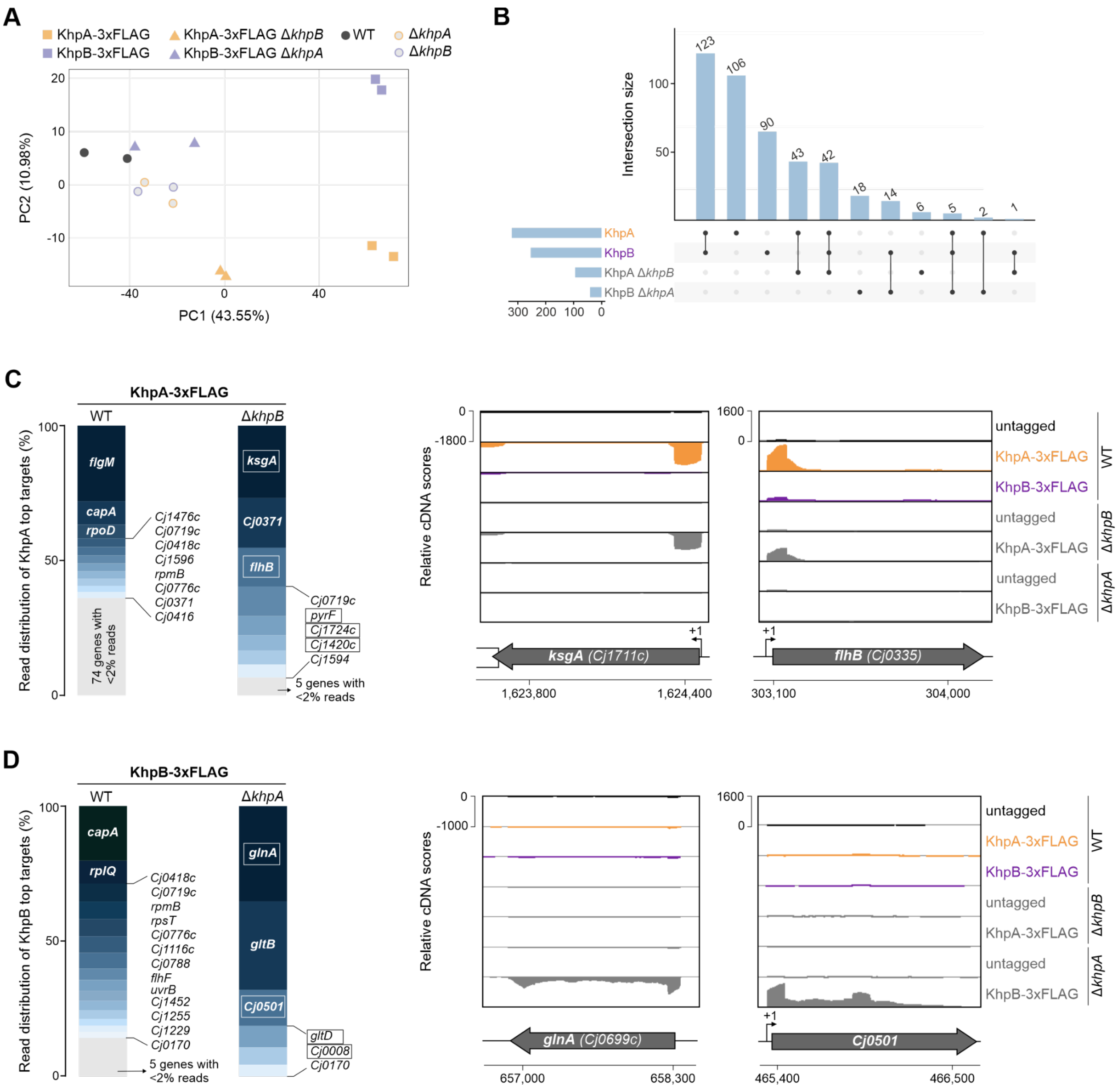


**Fig. S4: RNA targetome of KhpA and KhpB in the presence and absence of interacting partner.** **(A)** Principal component analysis (PCA) of 14 RIP-seq samples (corresponding to two replicates for each indicated strain), showing PC1 vs. PC2 in the WT and Δ partner background. Tagged libraries in WT condition are represented as squares, in Δ partner condition as triangles, and untagged libraries are shown in circles. **(B)** UpSet plot comparing the overlap of significantly enriched RNAs ($\log_2$ fold-change ≥1, *p*-adj ≤ 0.05, and average CPM ≥10 in tagged libraries) across the indicated RIP-seq datasets. Horizontal bars represent the number of co-purified RNAs, while vertical bars indicate intersection sizes. **(C, D)** (Left) Bar graphs showing the percentage of mapped reads for genes with >5-fold enrichment in KhpA-3xFLAG (C) and KhpB-3xFLAG (D) co-IP libraries in WT and Δ*khpB* (C) / Δ*khpA* (D) backgrounds. Gene names or locus names are indicated for each enriched gene. KhpA-specific targets enriched in Δ*khpB* (C) background are highlighted within a box. Unique targets enriched only in KhpB-3xFLAG Δ*khpA* background are highlighted within a box (D). Genes with less than 2% of mapped reads are grouped together and shown as the grey portion of each bar. (Right) Representative read coverage enrichment patterns in the 5' UTRs and/or CDSs of KhpA-specific top targets in the Δ*khpB* background (C) and top unique targets enriched only in KhpB-3xFLAG Δ*khpA* background (D). For the genome browser coverages, one representative biological replicate is shown. The transcriptional start site (TSS) is denoted as a black arrow.

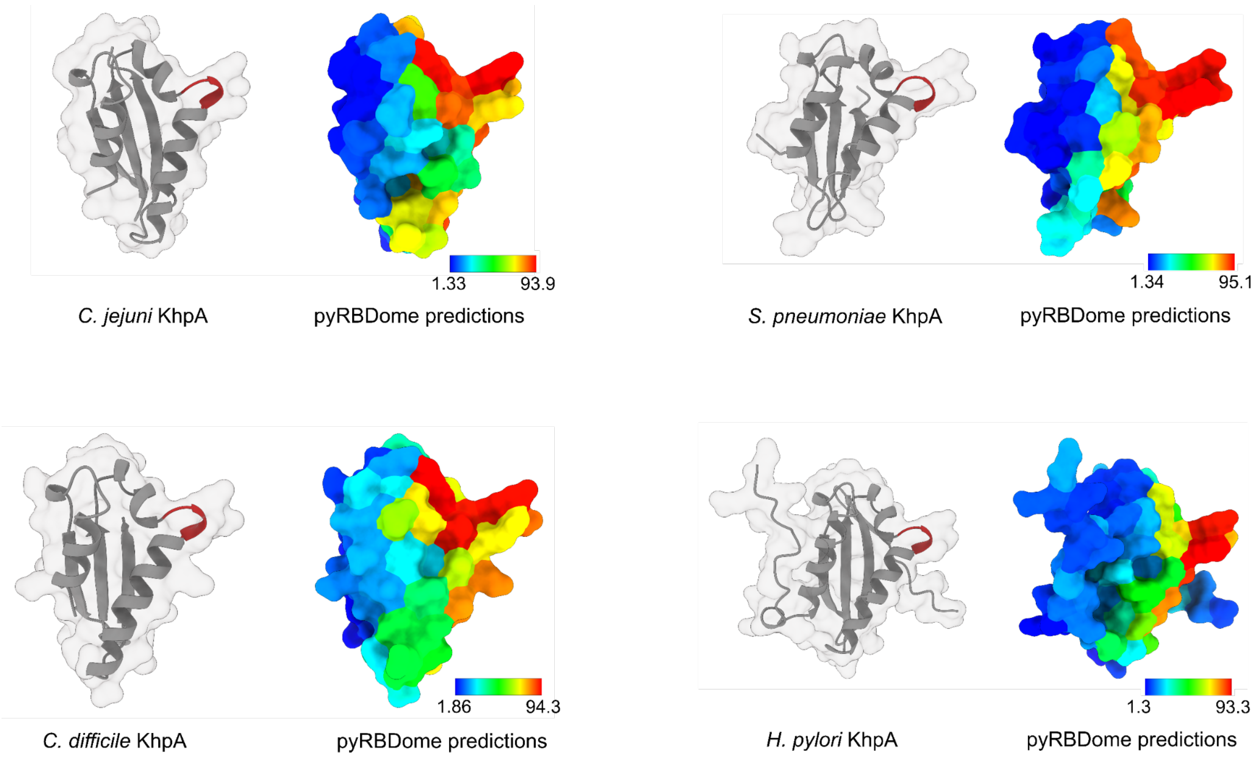


**Fig. S5: RNA-binding amino acids predicted by pyRBDome across KhpA homologs.** Structures retrieved from the UniProt database (left) and corresponding RNA-binding prediction outputs from the pyRBDome pipeline (right) are shown for KhpA homologs from *Campylobacter jejuni*, *Streptococcus pneumoniae*, *Clostridium difficile*, *and Helicobacter pylori*. The GXXG motif is highlighted in dark red for the respective KhpA homolog structure. For the pyRBDome (4) structures, cooler shades (blue) indicate amino acids with lower RNA-binding prediction scores, while warmer shades (red) denote amino acids with higher RNA-binding prediction scores.

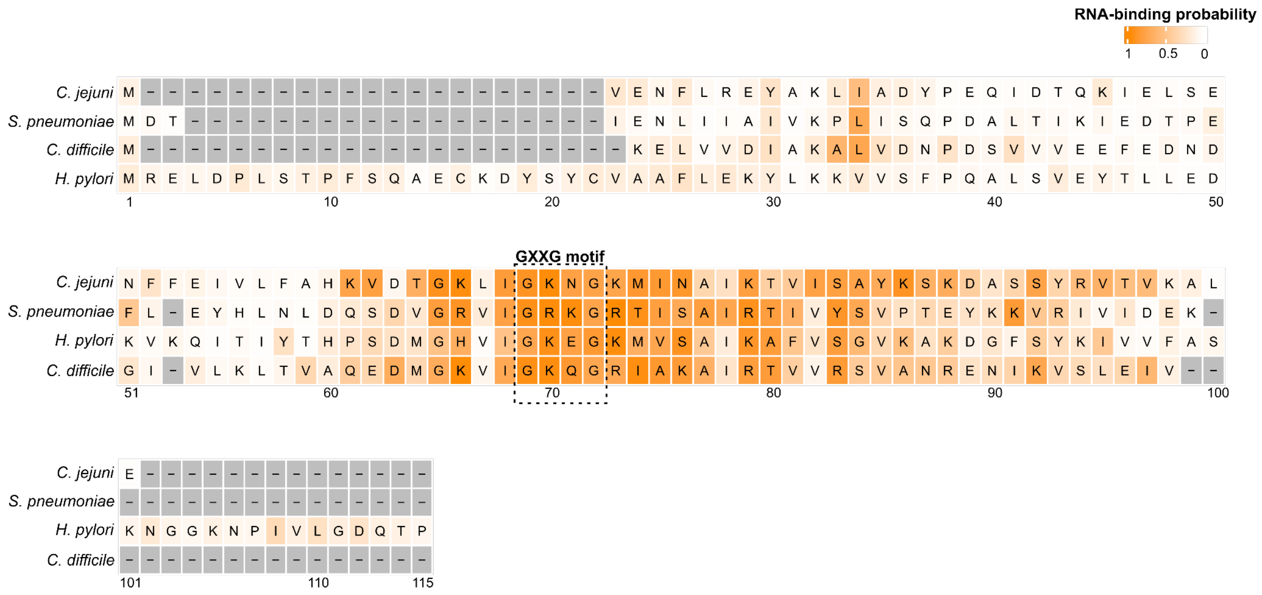


**Fig. S6: Heat map representation of predicted RNA-binding probabilities for individual amino acid residues across KhpA homologs.** Each row represents a KhpA homolog protein sequence from *Campylobacter jejuni*, *Streptococcus pneumoniae*, *Clostridium difficile*, *and Helicobacter pylori*, and each column corresponds to an aligned amino acid residue position. Intensity of orange represents the pyRBDome RNA-binding probabilities for each residue, with higher scores (darker shade) denoting a greater likelihood of RNA interaction. The GXXG motif of the KH domain is highlighted.

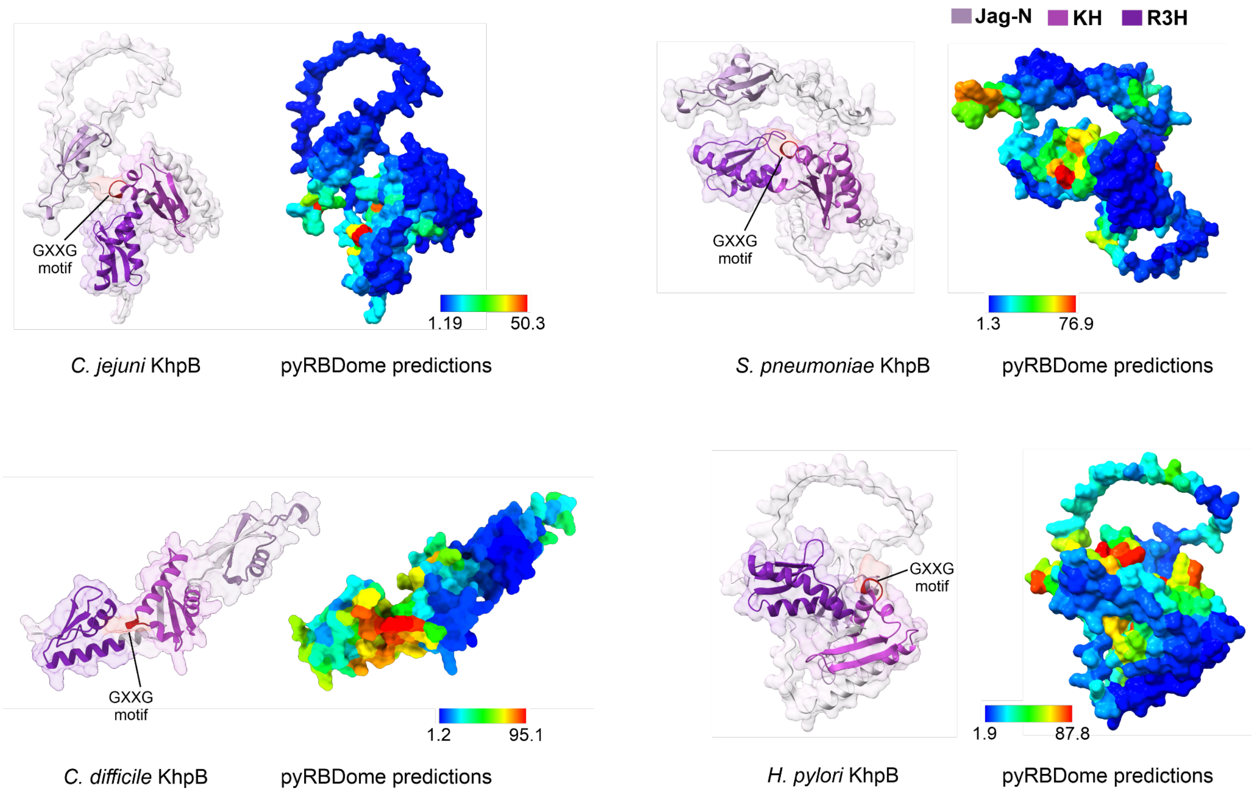


**Fig. S7: RNA-binding regions predicted by pyRBDome across KhpB homologs.** Structures retrieved from the UniProt database (left) and corresponding RNA-binding prediction outputs from the pyRBDome pipeline (right) are shown for KhpB homologs from *Campylobacter jejuni*, *Streptococcus pneumoniae*, *Clostridium difficile*, and *Helicobacter pylori*. Protein domains are colour coded according to the labels and the GXXG motif is highlighted in dark red within each KhpB homolog structure. In the pyRBDome panels, cooler shades (blue) indicate residues with lower predicted RNA-binding scores, while warmer shades (red) represent residues with higher predicted RNA-binding scores.

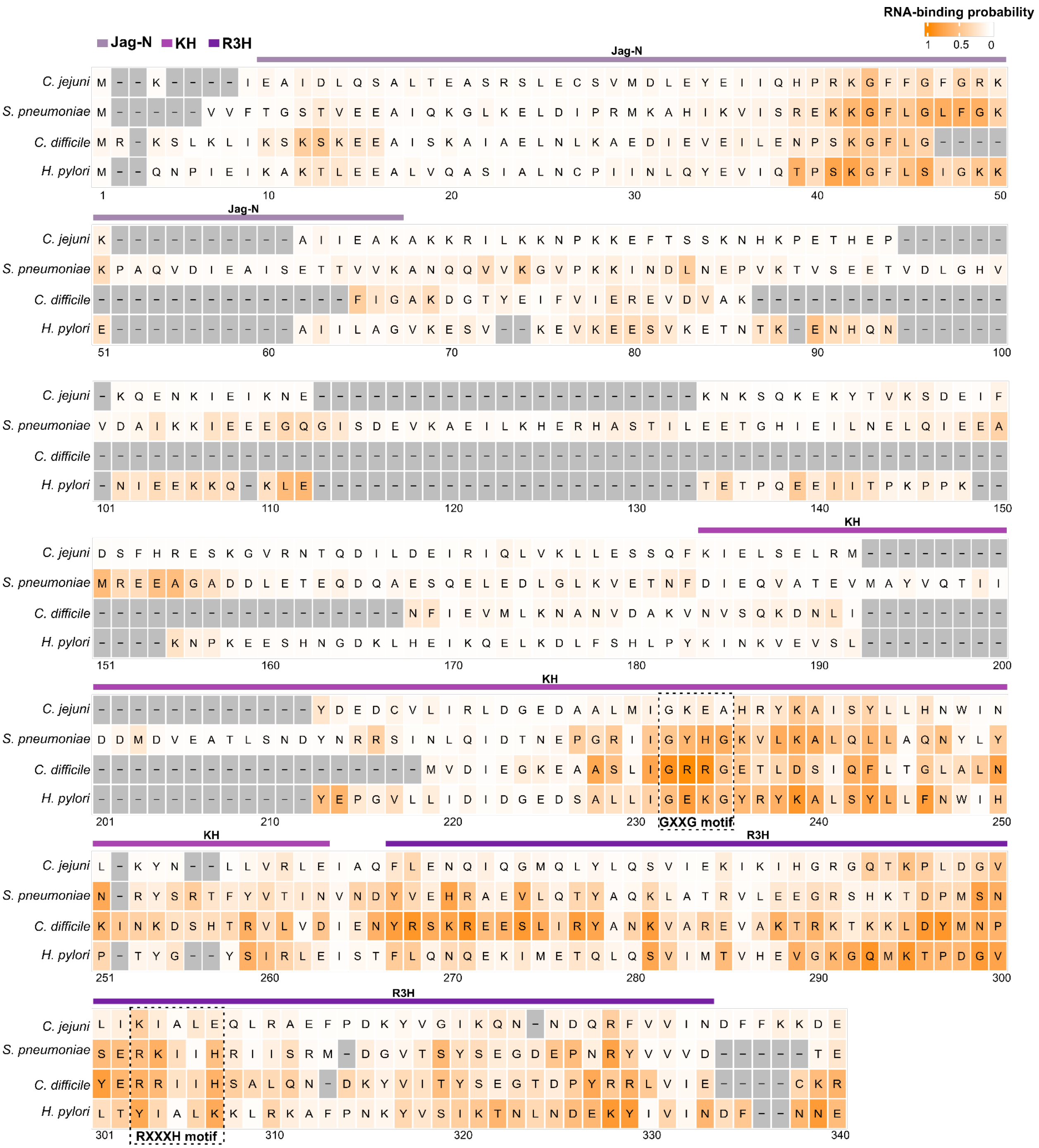


**Fig. S8: Heat map representation of predicted RNA-binding probabilities for individual amino acid residues across KhpB homologs.** Each row represents a KhpB homolog protein sequence from *Campylobacter jejuni*, *Streptococcus pneumoniae*, *Clostridium difficile*, *and Helicobacter pylori*, and each column corresponds to an aligned amino acid residue position. Intensity of orange represents the pyRBDome RNA-binding probabilities for each residue, with higher scores (darker shade) denoting greater likelihood of RNA interaction. Different domains and the RNA-binding motifs are highlighted.

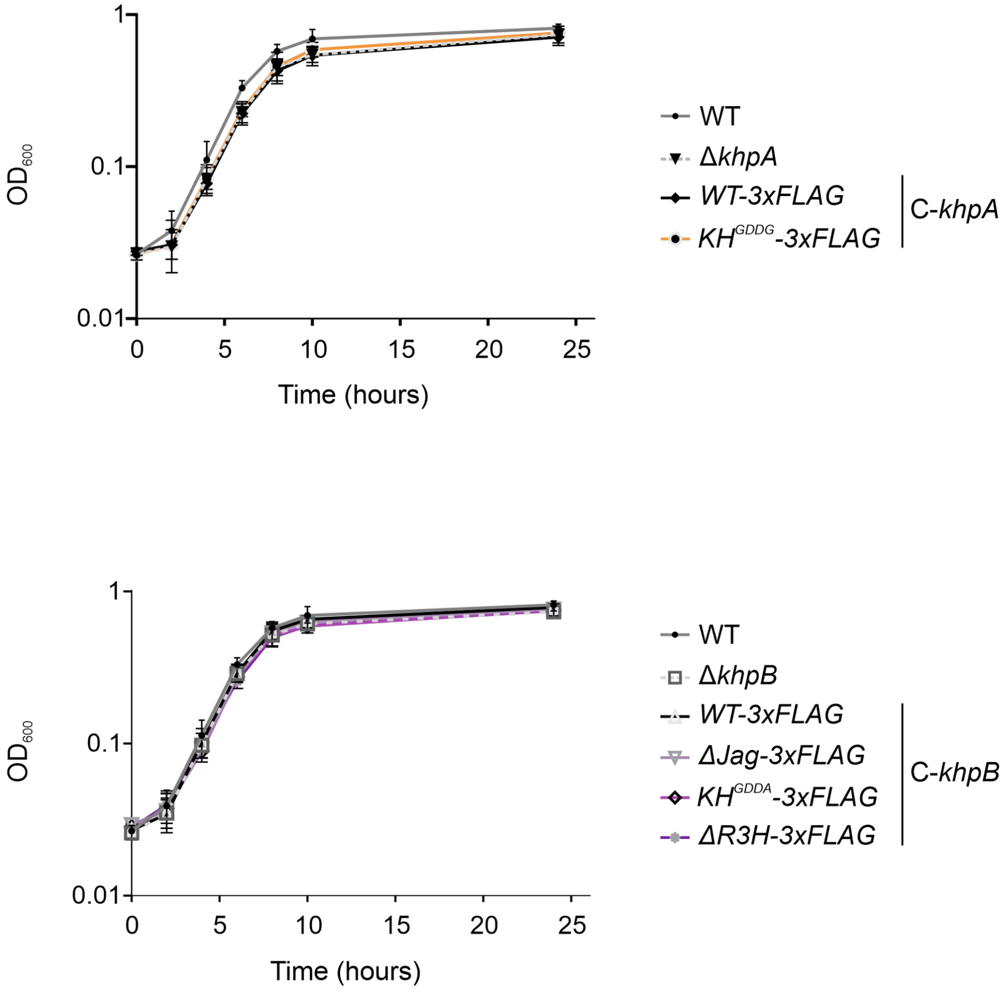


**Fig. S9: Growth curves of KhpA/B domain and point mutants.** Semi-log growth curves of *C. jejuni* NCTC11168 WT strain vs. 3xFLAG-tagged KhpA domain/point mutants (top) and KhpB domain/point mutants (bottom) over 24 h under standard growth conditions. Error bars indicate the standard deviation of three independent replicates.

**A**

| Pulled-down protein | Untagged background | Comparison control | |
|---|---|---|---|
| WT-3xFLAG | WT | - | C-*khpA* |
| $KH^{GDDG}$-3xFLAG | $KH^{GDDG}$ | WT-3xFLAG | C-*khpA* |
| WT-3xFLAG | WT | - | C-*khpB* |
| ΔJag-3xFLAG | ΔJag | WT-3xFLAG | C-*khpB* |
| $KH^{GDDA}$-3xFLAG | $KH^{GDDA}$ | WT-3xFLAG | C-*khpB* |
| ΔR3H-3xFLAG | ΔR3H | WT-3xFLAG | C-*khpB* |

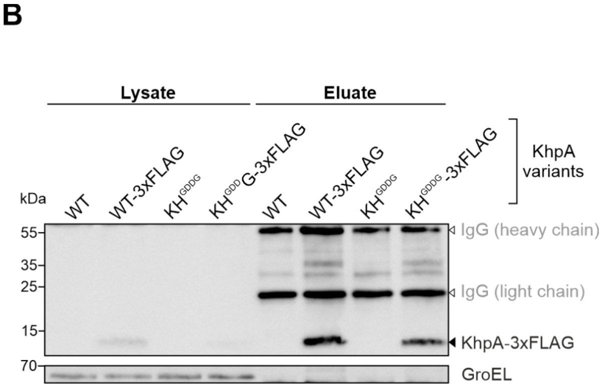


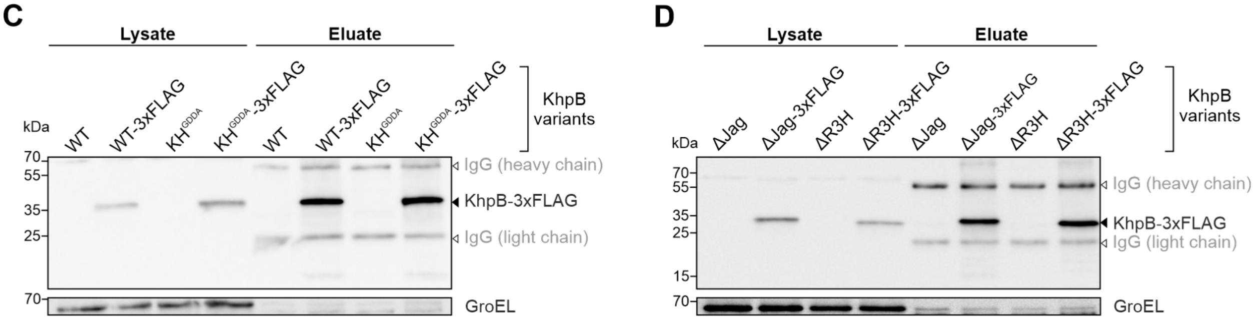


**Fig. S10: RIP-seq of WT and mutant domain variants of KhpA and KhpB. (A)** Table summarizing the FLAG-tagged pull-down mutants and corresponding untagged background strains used in RIP-seq experiments. Significantly enriched RNAs identified in each tagged mutant library are compared to the indicated WT controls. **(B-D)** Western blot analysis of immunoprecipitated proteins from *C. jejuni* lysates across different RIP-seq domain mutant experiments. FLAG-tagged proteins were detected using an anti-FLAG antibody. GroEL served as a loading control. One representative biological replicate out of two is shown.

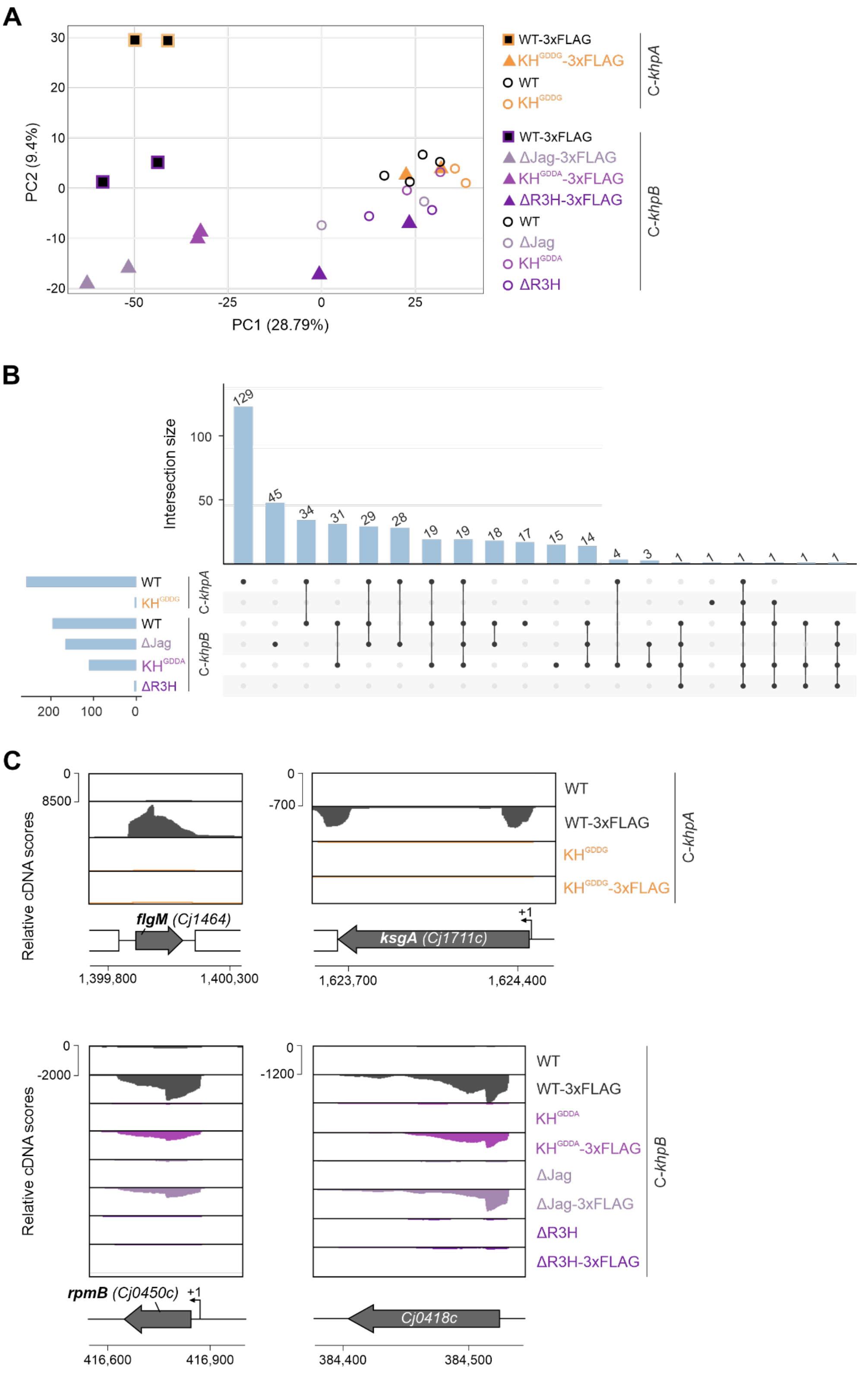


**Fig. S11: RNA targetome of WT and mutant domain variants of KhpA and KhpB. (A)** Principal component analysis (PCA) of 24 RIP-seq samples, showing PC1 vs. PC2 of WT and mutant domain variants of KhpA/B. Tagged libraries with WT domains are represented as squares, with mutant domains as triangles, and untagged libraries are shown in circles. Each of the 12 strains was tested in two biological replicates. **(B)** UpSet plot comparing overlaps of significantly enriched RNAs across the indicated RIP-seq datasets. Horizontal bars represent the number of co-purified RNAs, while vertical bars indicate intersection sizes. **(C)** Representative RIP-seq read coverage highlighting enrichment patterns in the 5' UTR and CDS regions of selected genes for WT vs. domain mutants of KhpA (top) and KhpB (below). One representative biological replicate is shown. The transcription start site (TSS) is denoted as a black arrow.

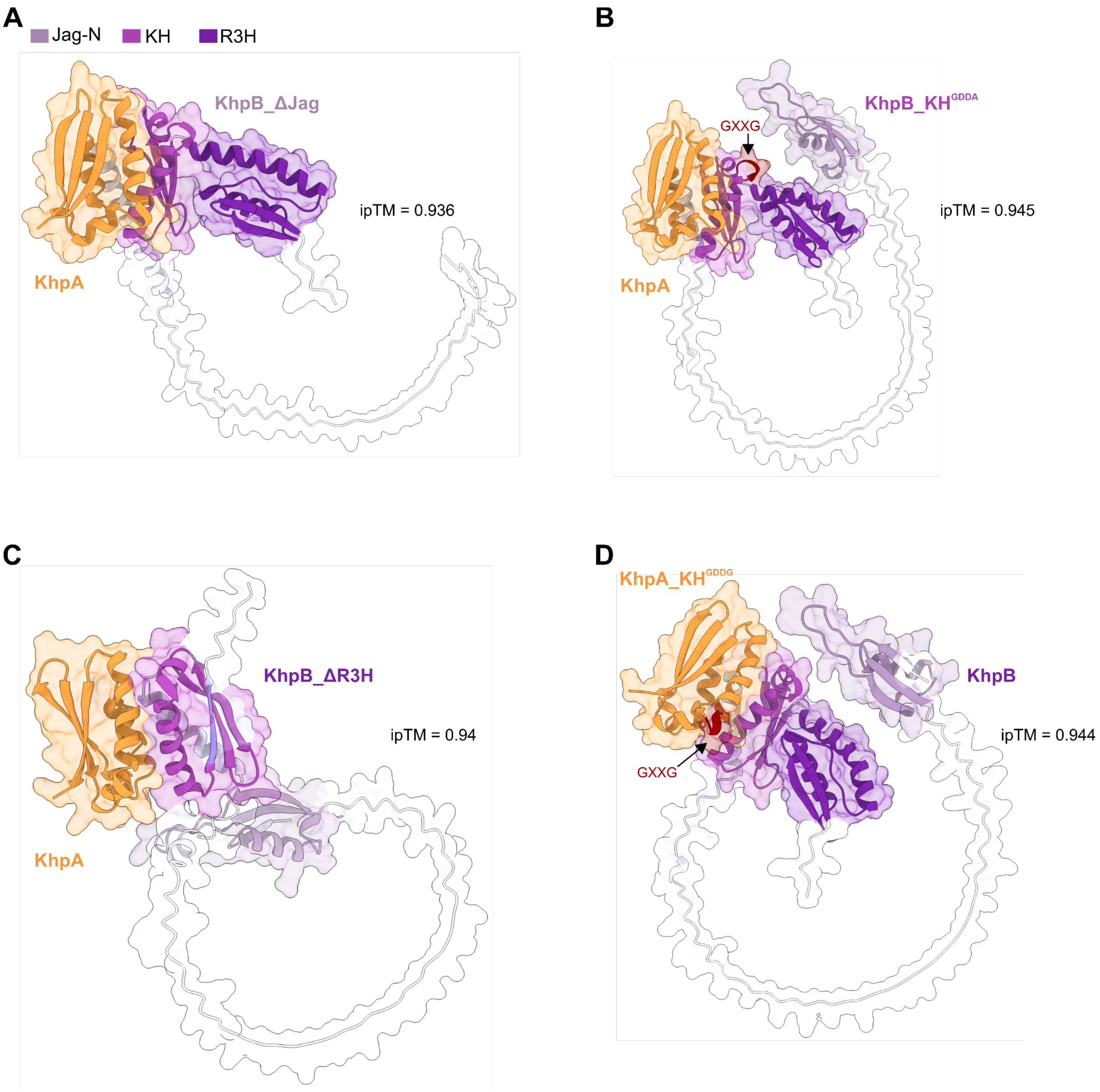


**Fig. S12: Predicted heterodimer structures of *C. jejuni* KhpA/B mutant domain variants. (A-C)** Predicted structural interactions between functional KhpA and various KhpB mutants: Jag-N-truncated KhpB (A), GDDA mutated KhpB (B), and R3H-truncated KhpB (C). **(D)** Predicted structural interactions between GDDG-mutated KhpA and functional KhpB. Dimers were modeled using AlphaFold2-multimer (5) and visualized with ChimeraX (6). Interface predicted template modelling (ipTM) for protein-protein interactions is depicted. Protein domains are colour coded according to the labels in (A).

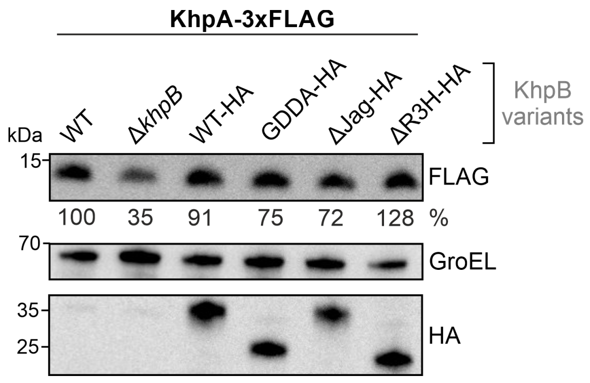


**Fig. S13: Protein expression levels of KhpA-3xFLAG in KhpB domain and point mutants.** Western blot analysis of KhpA-3xFLAG expression levels detected using an anti-FLAG antibody in the WT and mutant domain variants of KhpB-HA. Bands were quantified using ImageJ, normalized to GroEL, and percentages relative to KhpA-3xFLAG in *C. jejuni* WT are given. Blots are indicative of at least three independent replicates.

**A**

| Pulled-down protein | Untagged background control | Comparison control |
|---|---|---|
| KhpA-3xFLAG C-*khpB*_WT | C-*khpB*_WT | - |
| KhpA-3xFLAG C-*khpB*_ΔR3H | C-*khpB*_ΔR3H | KhpA-3xFLAG C-*khpB*_WT |
| KhpB-3xFLAG C-*khpA*_WT | C-*khpA*_WT | - |
| KhpB-3xFLAG C-*khpA*_KH$^{GDDG}$ | C-*khpA*_KH$^{GDDG}$ | KhpB-3xFLAG C-*khpA*_WT |

**B**

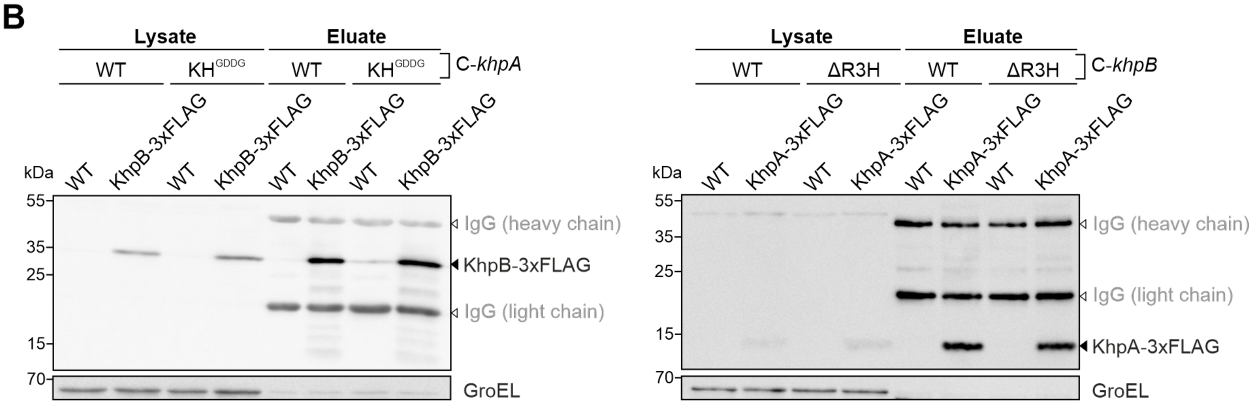


**Fig. S14: RIP-seq analyses of KhpA and KhpB in the WT and RBD mutant background of partner. (A)** Table summarizing the FLAG-tagged pull-down proteins in strains complemented with WT and RBD variants of partner protein (c-*khpA/B*), along with the corresponding untagged background strains complemented with the same partner protein variants (c-*khpA/B*), used in the RIP-seq experiments. Significantly e/nriched RNAs identified in each tagged mutant library were compared to the indicated WT controls. **(B)** Western blot analysis of immunoprecipitated proteins from *C. jejuni* lysates of KhpB-3xFLAG in *khpA*_WT and *khpA*_KH$^{GDDG}$ backgrounds (left) and KhpA-3xFLAG in *khpB*_WT and *khpB*_ΔR3H backgrounds (right). FLAG-tagged proteins were detected using an anti-FLAG antibody. GroEL served as a loading control. One representative biological replicate out of two is shown.

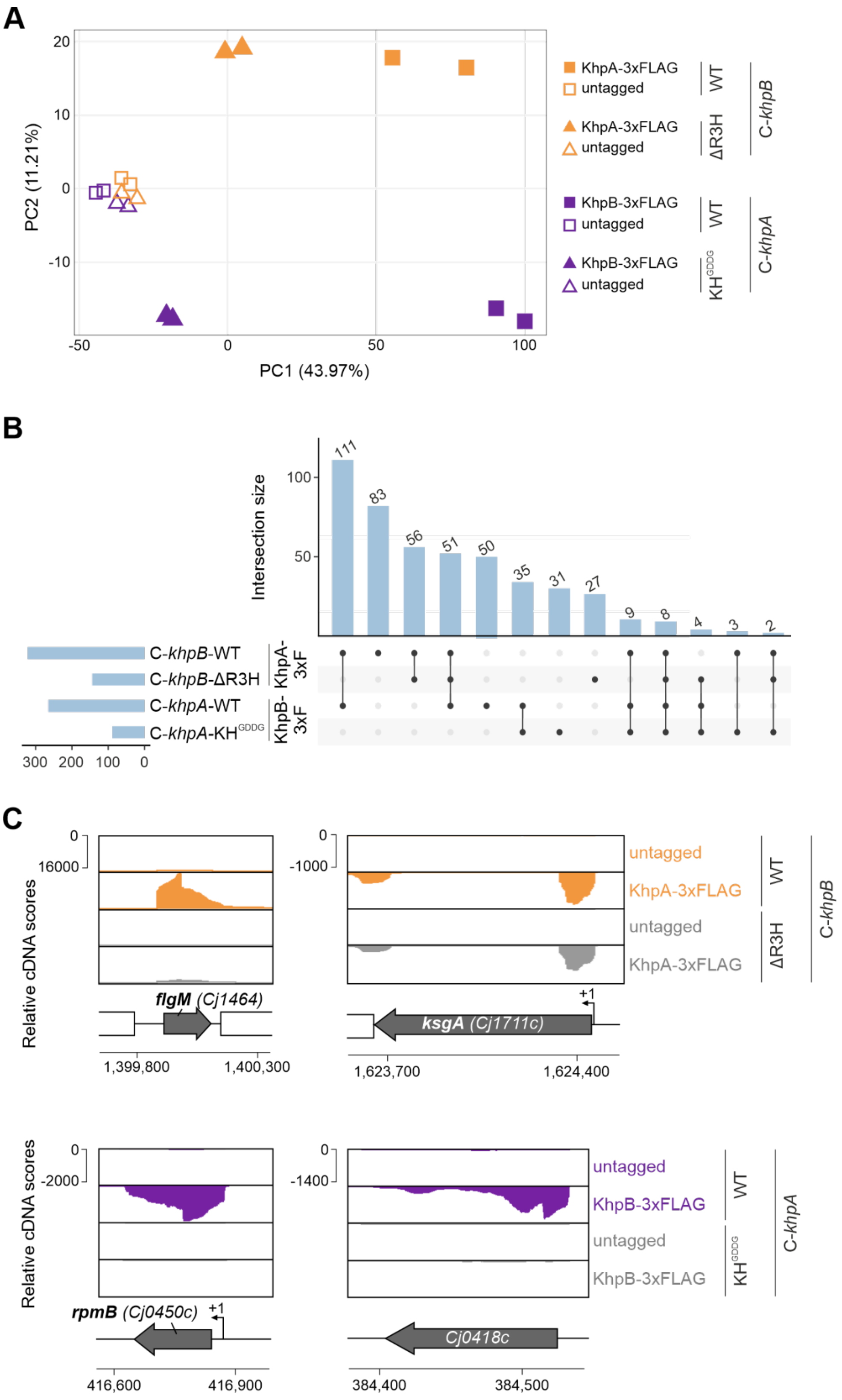
A
PC1 (43.97%)
PC2 (11.21%)
KhpA-3xFLAG
untagged
WT
ΔR3H
C-khpB
KhpB-3xFLAG
KH^GDDG
C-khpA
B
Intersection size
C-khpB-WT
C-khpB-ΔR3H
C-khpA-WT
C-khpA-KH^GDDG
KhpA-3xF
KhpB-3xF
111 83 56 51 50 35 31 27 9 8 4 3 2
C
Relative cDNA scores
flgM (Cj1464)
ksgA (Cj1711c)
rpmB (Cj0450c)
Cj0418c
+1
1,399,800
1,400,300
1,623,700
1,624,400
416,600
416,900
384,400
384,500

**Fig. S15: RNA targetome of KhpA and KhpB in the WT and RBD mutant background of the partner. (A)** Principal component analysis (PCA) of 16 RIP-seq samples, showing PC1 vs. PC2 of KhpA and KhpB in WT and RBD mutant background of partner. Each condition includes two biological replicates. **(B)** UpSet plot comparing overlaps of significantly enriched RNAs (≥2-fold enrichment) across the indicated RIP-seq datasets. Horizontal bars represent the number of co-purified RNAs, while vertical bars indicate intersection sizes. **(C)** Representative RIP-seq read coverage highlighting enrichment patterns in the 5' UTR and CDS of selected genes for KhpA in WT and $khpB_{\Delta\mathrm{R3H}}$ backgrounds (top) and KhpB in WT and $khpAB_{\mathrm{GDDG}}$ backgrounds (below). One representative biological replicate is shown.

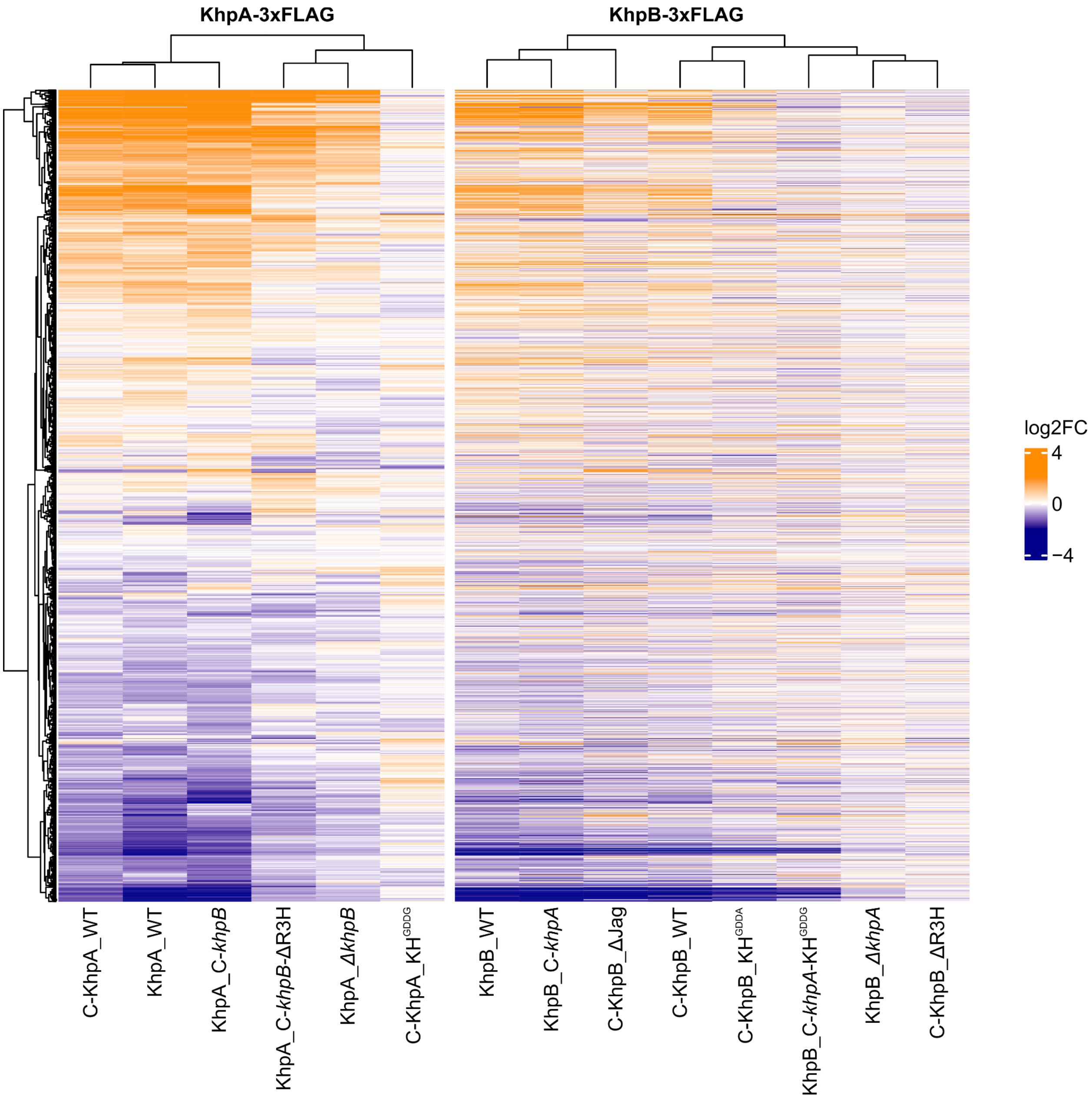


**Fig. S16: Hierarchical clustering heatmap of all RIP-seq datasets.** The heatmap illustrates RNA-binding profiles of FLAG-tagged strains, displaying $log_2$ fold changes ($log_2FC$) calculated from DESeq2 analysis comparing each FLAG-tagged strain to its respective untagged control. Rows represent individual transcripts, while columns represent FLAG-tagged pull-down samples. Colours indicate $log_2FC$ values, and hierarchical clustering reveals patterns of similarities in RNA-binding profiles across samples.

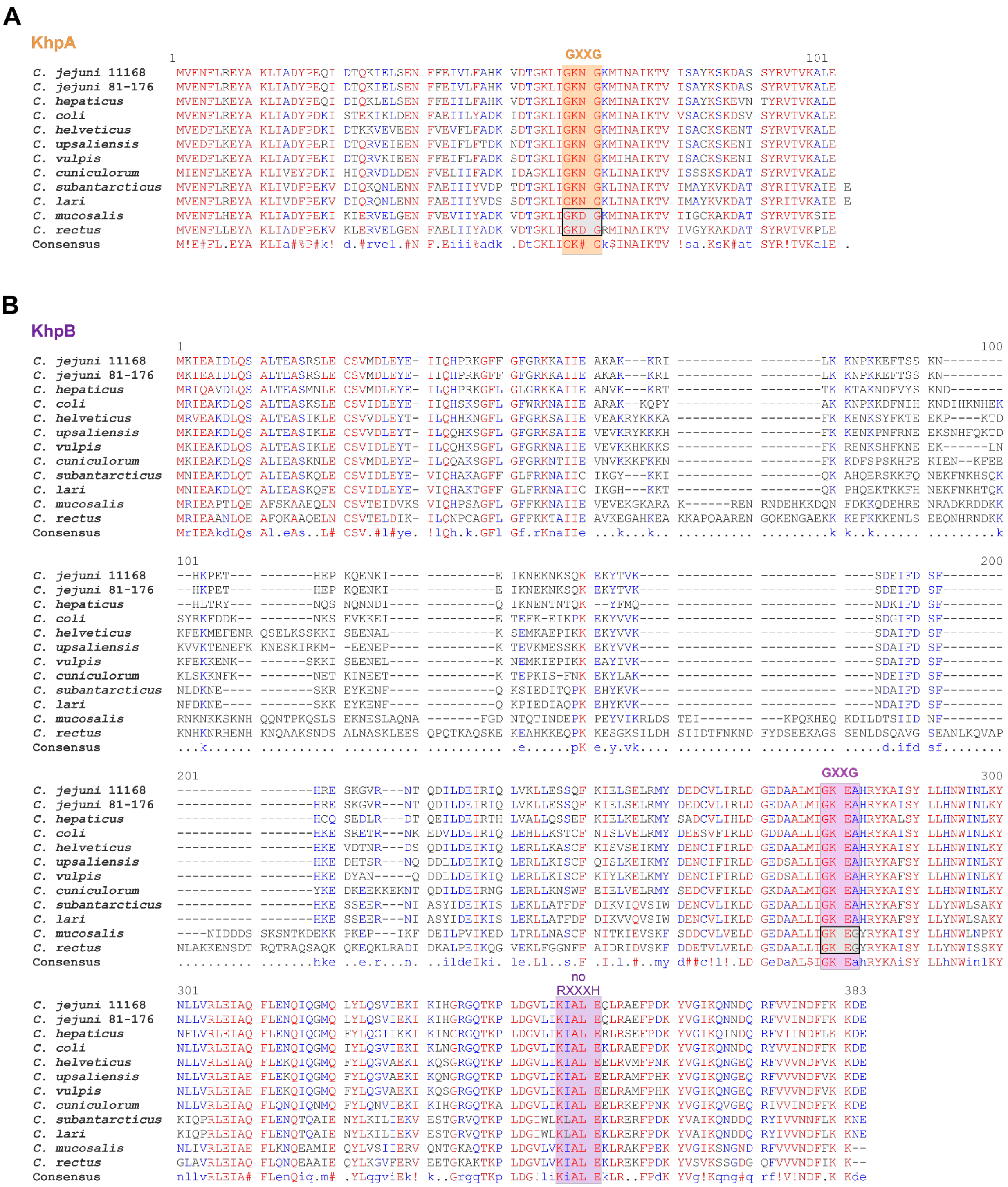


**Fig. S17: Multiple sequence alignment of the GXXG motif and the R3H motif across *Campylobacter* species. (A-B)** Alignment of protein sequences of KhpA (A) and KhpB (B) from selected *Campylobacter* species using Multalin (1). The GXXG motif of the KH domain and the RXXXH motif of the R3H domain are highlighted. The GXXG motifs of the *Campylobacter* species that show co-variation between KhpA and KhpB are highlighted in gray.

### Supplementary References


1. F. Corpet, Multiple sequence alignment with hierarchical clustering. *Nucleic Acids Res.* **16**, 10881–10890 (1988).
2. S. L. Dove, J. K. Joung, A. Hochschild, Activation of prokaryotic transcription through arbitrary protein-protein contacts. *Nature* **386**, 627–630 (1997).
3. K. E. Berry, A. Hochschild, A bacterial three-hybrid assay detects *Escherichia coli* Hfq-sRNA interactions in vivo. *Nucleic Acids Res.* **46**, e12 (2018).
4. L.-C. Chu, *et al.*, pyRBDome: a comprehensive computational platform for enhancing RNA-binding proteome data. *Life Sci. Alliance* **7** (2024).
5. J. Jumper, *et al.*, Highly accurate protein structure prediction with AlphaFold. *Nature* **596**, 583–589 (2021).
6. E. F. Pettersen, *et al.*, UCSF ChimeraX: structure visualization for researchers, educators, and developers. *Protein Sci.* **30**, 70–82 (2021).
